# An Unconventional Melanin Biosynthetic Pathway in *Ustilago maydis*

**DOI:** 10.1101/2019.12.28.889758

**Authors:** Esmeralda Z. Reyes-Fernández, Yi-Ming Shi, Peter Grün, Helge B. Bode, Michael Bölker

## Abstract

*Ustilago maydis* is a phytopathogenic fungus responsible for corn smut disease. Although it is a very well established model organism for the study of plant-microbe interactions, its biosynthetic potential has not been totally explored. By analyzing *U. maydis* genome, we identified a biosynthetic gene cluster whose activation led to the production of a black melanin pigment. Single deletion mutants of the cluster genes revealed that five encoded enzymes are required for the accumulation of the black pigment, including three polyketide synthases (*pks3, pks4* and *pks5*), a cytochrome P450 monooxygenase (*cyp4*) and a protein with similarity to versicolorin B-synthase (*vbs1*). Moreover, metabolic profiles of the mutants defective for *pks3* and *pks4* indicated that the products of these genes catalyze together the first step in the melanin biosynthetic pathway since none of the mutants accumulated any melanin or intermediate products. Mutants deleted for *pks5* produced orsellinic acid (OA) and triacetic acid lactone (TAL), suggesting that both products are produced by Pks3 and Pks4. It might thus demonstrate that Pks5 plays a role in a reaction downstream of that catalyzed by Pks3 and Pks4. OA and TAL were also found in extracts of a *cyp4* deletion mutant along with several heterodimers of TAL and Pks5-derived orsellinic aldehyde compounds. According to their phenotypes and the intermediate products isolated from these strains, Cyp4 and Vbs1 seem to be involved in reactions downstream of Pks5. Our findings suggest that *U. maydis* synthesizes a new melanin based on coumarin and pyran-2-one intermediates, while most fungal melanins are derived from 1,8-dihydroxynaphthalene (DHN) or L-3,4-dihydroxyphenylalanine (L-DOPA). Along with these observations, this work also provides an insight into the mechanisms of polyketide synthases in this filamentous fungus.

**IMPORTANCE:** *Ustilago maydis* represents one of the major threats for maize plants since it is responsible for corn smut disease, which generates considerable economical losses around the world. Therefore, contributing to a better understanding of the biochemistry of defense mechanisms used by *U. maydis* to protect itself against harsh environments, as the synthesis of melanin, could provide improved biological tools for tackling the problem and protect the crops. In addition, the fact that this fungus synthesizes melanin in a very unique way, requiring more than one polyketide synthase for producing this secondary metabolite, gives a different perspective on the complexity of these multimodular enzymes and their evolution in the fungal kingdom.

## INTRODUCTION

Secondary metabolites (SMs) produced by bacteria, plants and fungi are a vast group of small molecules with fascinating bioactivities that play non-essential roles in growth, development and reproduction of a living organism. Genes involved in the biosynthesis of fungal SMs are generally fast-evolving, often clustered and co-regulated through common transcription factors or at the level of chromatin organization (1). Despite the potential versatility of fungi in producing SMs, their biosynthetic pathways often have not been well explored since the biosynthetic gene clusters are silent under standard laboratory conditions (2). Methods to activate silent biosynthetic pathways are thus of major interest (3, 4). Examples of SM that can be observed by our naked eye are pigments, which absorb damaging radiation and thus protect the organism against DNA damage and oxidative stress (5). Among those pigments, melanin plays an essential role in many organisms, including fungi (6). Phenolic and/or indolic monomers are prone to polymerize forming negatively charged, hydrophobic melanins (7), which contribute to the ability of fungi to survive in harsh environments. Besides, it plays a role in pathogenesis (8).

In fungi, two main melanin biosynthetic pathways have been described, 1,8-dihydroxynaphthalene (DHN) and L-3,4-dihydroxyphenylalanine (L-DOPA), with the DHN pathway being more widespread (9, 10).

*Ustilago maydis*, a hemibiotrophic basidiomycete, is the causative agent of maize (*Zea mays*) smut disease. The disease cycle is initiated by the fusion of compatible haploid cells. The resulting dikaryon is filamentous, grows in close contact with the plant and is able to sense signals from the leaf surface that trigger the formation of appressoria (11). The plasma membrane of the plant cell invaginates and tightly surrounds the invading hypha (12). An interaction zone develops between the plant and fungal membranes that is characterized by fungal deposits produced by exocytosis. At later stages, proliferation also occurs intercellularly and the dikaryotic mycelium grows towards bundle sheets (13). Proliferation is followed by karyogamy and sporogenesis, where hyphal sections fragment, round up and differentiate into heavily melanized diploid teliospores (14).

The first studies of melanin in *U. maydis* suggested that catechol was the precursor for its biosynthesis since this compound was found in ethanol extracts from teliospores (15). A recent study showed that two PKSs (Pks1 and Pks2) and a laccase (Lac1) are involved in the melanization process of teliospores during plant infection (16). That work gave a preliminary idea about the role of Pks1 and Pks2, but the biosynthetic pathway of melanin production during spore formation is yet to be characterized.

Here, we describe the identification and characterization of a silent PKS based biosynthetic gene cluster in *U. maydis* involved in alternative melanin production. Expression of the pathway-specific transcription factor *mtf1* activated at least 12 genes encoding three polyketide synthases (*pks3, pks4* and *pks5*), a cytochrome P450 (*cyp4*) and a versicolorin B-synthase like protein (*vbs1*), among others. Activation of this gene cluster resulted in the production of black melanin pigment. By generating deletion mutants and analysis of their metabolic profiles by LC-MS, we were able to show that mutants defective in either one of the PKS-encoding genes *pks3, pks4, pks5* or the cytochrome P450 gene *cyp4* did not accumulate melanin, thus indicating a crucial role of the corresponding enzymes for pigment biosynthesis. Furthermore, single and combined expression of these PKS genes revealed that *pks3* and *pks4* are together required for generating orsellinic acid (OA) and triacetic acid lactone (TAL). These intermediates undergo further enzymatic modifications carried out by Pks5, Cyp4, Vbs1 and other proteins to yield melanin. These data demonstrate that unlike any other fungi, *U. maydis* possesses a unique mechanism for melanin synthesis that requires the collaboration of at least three PKSs, a cytochrome P450 and a versicolorin B-synthase like enzyme.

## RESULTS

### Identification of a co-regulated PKS biosynthetic gene cluster in *U. maydis*

Preliminary work has shown that *U. maydis* is capable of synthesizing diverse secondary metabolites (17, 18). In order to unveil more secondary metabolites in *U. maydis*, we decided to seek for potential biosynthetic gene clusters encoding PKSs, typical backbone enzymes involved in secondary metabolism (19). By analyzing the *U. maydis* genome database (MUMDB), we noticed the presence of three genes encoding PKS located in close proximity to each other on the telomeric region of chromosome 12 **(Fig. 1A and B)**. We termed these genes as *pks3 (UMAG_04105), pks4* (*UMAG_04097)* and *pks5 (UMAG_04095)*. Subsequently, we identified in the neighboring area of *pks3*-5 two genes coding for putative transcription factors, which were termed *mtf1 (UMAG_04101)* and *mtf2 (UMAG_11110*). This suggests that *pks3, pks4* and *pks5* could be part of a silent biosynthetic gene cluster, whose regulation was potentially controlled by either *mtf1* or *mtf2.* To test this hypothesis, we generated two strains, in which each transcription factor (*mtf1* or *mtf2*) was expressed under the control of the arabinose-inducible P*crg* promoter (P*crg*::*mtf1* and P*crg*::*mtf2*) (20). Afterward, we checked by Northern blot, which genes were regulated by which transcription factor **(Fig. S1A).** Transcription of the genes *mtf1* and *mtf2* was detectable in the selected transformants only under inducing conditions (I) and completely absent under non-inducing conditions (R) **(Fig. S1A)**. Notably, when the P*crg*::*mtf1* strain was grown under inducing conditions, *pks3, pks4* and *pks5* were substantially transcribed. Likewise, *UMAG_04096* (*orf1*), *UMAG_11111* (*aox1*), *UMAG_11112* (*vbs1*), *UMAG_04104* (*orf4*), *UMAG_04106* (*omt1*), *UMAG_04107* (*pmo1*), *UMAG_12253* (*orf5*), *UMAG_04109* (*cyp4*) and *UMAG_11113* (*deh1*) genes were expressed upon induction of *mtf1.* Conversely, no transcript of these genes was detected under non-inducing conditions. On the other hand, over-expression of *mtf2* only up-regulated the gene *UMAG_04098* (*orf2*), thus indicating that activation of *mtf2* has no influence in the expression of the other genes **(Fig. S1A)**. This result allowed us to establish the boundaries of the potential biosynthetic gene cluster **(Fig. 1A)**. The cluster expands from the most-upstream gene *pks5* until the most-downstream gene *deh1* with a total length of 45.1 kb, containing three genes in between (*orf2, mtf2* and *orf3*) whose expression is not controlled by *mtf1.* A detailed description of the cluster genes is shown in **Fig. 1B**. Together, the present findings confirm that this gene cluster **(Fig. 1A)** remains silent when *U. maydis* is grown under normal laboratory conditions and that expression of *mtf1* is sufficient to activate the cluster genes.

**Fig. 1.**
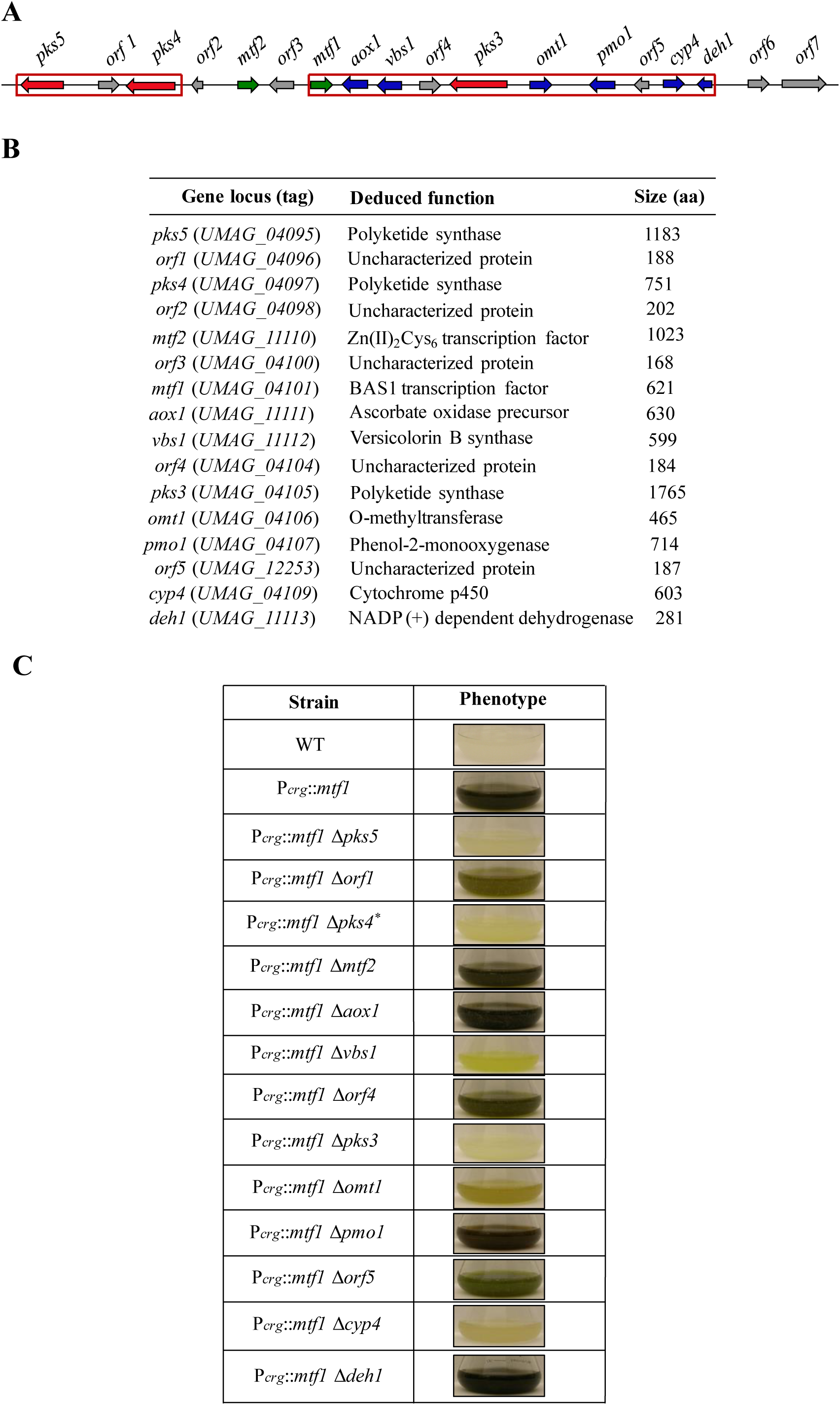
Induction of a large polyketide synthase gene cluster results in the production of a black-greenish pigment. **(A)** The identified PKS-cluster consists of at least 12 co-regulated genes. Among these are three genes encoding polyketide synthases (*pks3, pks4*, and *pks5*) and several other genes characteristic of secondary metabolism. Protein-encoding genes belonging to different categories are highlighted as follows: red, backbone enzyme; blue, tailoring enzyme and gray, other. **(B)** Gene locus, deduced function and size of the protein-coding genes located within the melanin gene cluster. **(C)** Phenotype of the melanin cluster deletion mutants. Single-deletion mutants of the melanin cluster were generated in the P*crg*::*mtf1* background of the MB215 strain. Photographs of the cultures were taken after 96 h of growth at 28 °C in inducing medium.

We also noticed that in liquid media, prolonged expression of the PKS gene cluster in strain P*crg::mtf1* triggered the production of a black-greenish pigment, which was clearly observed after 24 h of induction **(Fig. S1B)**. In contrast, no effect was recorded in the P*crg::mtf2* and wild-type strains **(Fig. S1B)**. Similarly, dark colonies were observed only for the P*crg::mtf1* strain when wild-type, P*crg::mtf1* and P*crg::mtf2* cells were spotted in serial dilutions on inducing agar medium (data not shown).

### Pigmentation phenotype of gene deletions

In order to determine the biological role of this biosynthetic gene cluster for pigment production, we generated single deletion mutants of all 12 genes regulated by *mtf1.* The coding sequences were replaced by a selectable marker gene in the genetic background of strain P*crg*::*mtf1.* Transformants were analyzed by PCR and Southern blot. In the case of *pks4*, we succeeded in replacing the gene but Southern blot always showed that both the expected band of the deletion mutant and the wild-type band were still detectable. For that reason, we suspected that *pks4* could be duplicated in the *U. maydis* genome. Only after deleting the duplicated copy of *pks4* in the P*crg*::*mtf1* Δ*pks4* strain, the signal corresponding to the *pks4* wild-type band was absent in the Southern blot. We termed the strain containing the double mutant of *pks4* as P*crg*::*mtf1* Δ*pks4*\*. In the FB1-derived strain P*crg*::*mtf1* (FB1) deletion of a single copy of *pks4* was sufficient to result in loss of melanin production. This indicates that duplication of *pks4* is characteristic of the genetic background of strain MB215 but obviously has not occurred in FB1. For the other two genes located upstream of *pks4, orf1* and *pks5*, we encountered the same problems while trying to delete them in the P*crg*::*mtf1* (MB215). Therefore, we generated the corresponding deletions also in strain P*crg*::*mtf1* (FB1).

As shown in **Fig. 1C**, deletion mutants of either *pks3, pks4, pks5* and *cyp4*, a gene encoding a cytochrome P450, were significantly affected in their phenotype and were unable to synthesize the melanin-like pigment **(Fig. S2)**. Deletion of *vbs1*, a gene that codes for a protein with sequence similarity to versicolorin B synthase (VBS) from *Aspergillus parasiticus* (21), resulted in a yellowish phenotype **(Fig. 1C).** Strain P*crg*::*mtf1* Δ*omt1* accumulated an orange-yellowish pigment, which was in appearance darker than that produced by the P*crg*::*mtf1 Δvbs1*. Strains deleted for *aox1* (ascorbate oxidase), *pmo1* (phenol-2-monooxygenase), or one of the remaining ORFs (*orf1*-*orf5*) showed no detectable phenotype **(Fig. 1C)**. In summary, disruption mutants of *pks3, pks4, pks5* and *cyp4* abolished synthesis of the melanin-like pigment, thus indicating the crucial role of these genes in the biosynthetic pathway. Although neither deletion of *vbs1* nor *omt1* produced a colorless phenotype, the participation of the encoded enzymes in melanin production was suggested by their yellowish and dark yellowish phenotypes, respectively. Only small changes in pigmentation were observed in the P*crg*::*mtf1* Δ*orf1*, P*crg*::*mtf1* Δ*aox1*, P*crg*::*mtf1* Δ*orf4*, P*crg*::*mtf1* Δ*pmo1*, P*crg*::*mtf1* Δ*orf5* and P*crg*::*mtf1* Δ*deh1* strains if compared with the reference, suggesting a minor involvement of these gene products in the production of the melanin-like pigment.

### Metabolic profiles and chemical identification of compounds produced by the mutant strains of the melanin-like gene cluster

By mutational analysis of all the genes that were up-regulated upon activation of *mtf1*, we gained an insight into the enzymes that are involved in the biosynthetic pathway of melanin-like pigment in *U. maydis*. Intending to have a deeper understanding of the sequential steps of the pathway, we decided to analyze the cell pellet extracts of all the single mutants by LC-MS. Notably, analysis extracts of P*crg*::*mtf1* Δ*pks3* did not reveal any detectable compound **(Fig. 2)**, suggesting that Pks3 is essential for the biosynthesis of the pigment, as well as indicating its crucial role at first stages in the metabolic pathway. For the mutant of *pks4*, we analyzed the metabolic profiles of the single (P*crg*::*mtf1* Δ*pks4*) and double (P*crg*::*mtf1* Δ*pks4\**) deletion strains **(Fig. 2** and **Fig. S2B)**. Deletion of both copies of *pks4* in the P*crg*::*mtf1* background strain produced no compounds, while the extracts of the P*crg*::*mtf1* Δ*pks4* exhibited a similar metabolic profile as the parental strain P*crg*::*mtf1* **(Fig. S2B)**. To exclude the possibility that the colorless phenotype of P*crg*::*mtf1* Δ*pks4\** could be due to a defect in the expression of those genes whose deletion triggered a similar phenotype (*pks3, pks5* and *cyp4*), we analyzed their transcripts levels by Northern blot **(Fig. S2A)**. As expected, the expression of *pks3, pks5* and *cyp4* remained unaffected, thus suggesting that the phenotype and metabolic profile of P*crg*::*mtf1* Δ*pks4\** was solely attributed to the deletion of both copies of *pks4*. On the other hand, the LC-MS profile of the P*crg*::*mtf1* Δ*pks5* strain generated two peaks that corresponded to compounds **1** and **3 (Fig. 2)**. Those compounds were isolated and their structures elucidated by NMR. Compound **1** (*m*/*z* =279 [M+H]^+^) was identified as orsellinic acid (OA), while compound **3** (*m*/*z* =127 [M+H]^+^) was identified as 4-hydroxy-6-methyl-2-pyrone, also known as triacetic acid lactone (TAL). Likewise, the extracts of P*crg*::*mtf1* Δ*cyp4* indicated the presence of the same compounds, OA (**1**) and TAL (**3**), although the amount of TAL was much lower compared to the P*crg*::*mtf1* Δ*pks5* strain **(Fig. 2)**. Along with OA and TAL, extracts of P*crg*::*mtf1* Δ*cyp4* also produced the compounds **2** (*m*/*z* =153 [M+H]^+^), corresponding to 2,4-dihydroxy-6-methyl-benzaldehyde (orsellinic aldehyde), and **4**/**5** (*m*/*z* =263 [M+H]^+^), identified as 7-hydroxy-3-(3-hydroxybutanoyl)-5-methylcoumarin and 3-(2,4-dihydroxy-6-methylbenzyl)-4-hydroxy-6-methyl-2H-pyran-2-one, respectively. All these data suggest that Pks3 and Pks4 participate in the first metabolic step, followed by the reactions catalyzed by Pks5 and then by Cyp4. Subsequently, we analyzed the strain P*crg*::*mtf1* Δ*vbs1*, which accumulated a yellowish pigment **(Fig. 1C)**. The LC-MS analysis of the strain presented two major peaks (**6** and **7**), which were analogous to the compounds 7-hydroxy-3-(3-hydroxybutanoyl)-5-hydroxymethylcoumarin and 3-(2,4-dihydroxy-6-(hydroxymethyl)benzyl)-4-hydroxy-6-methyl-2H-pyran-2-one, respectively **(Fig. 2)**. Based on the chemical structure of **6** and **7** elucidated by NMR **(Fig. S5-S14 and Table S5)** and the phenotype of P*crg*::*mtf1* Δ*vbs1*, it can be inferred that Vbs1 contributes to the biosynthesis of the actual melanin-like pigment by reactions downstream from the one catalyzed by Cyp4. In line with this observation, the double deletion of *vbs1* and *cyp4* produced the same compounds found in the extracts of the single mutant of *cyp4*, thus indicating that Vbs1 indeed acts downstream of Cyp4 (data not shown). Finally, the chromatograms of the strains P*crg*::*mtf1* Δ*aox1*, P*crg*::*mtf1* Δ*orf4*, P*crg*::*mtf1* Δ*omt1*, P*crg*::*mtf1* Δ*pmo1*, P*crg*::*mtf1* Δ*orf5*, and P*crg*::*mtf1* Δ*deh1* mimicked the profile of the reference strain (data not shown). The evidence presented in this section suggests that there are sequential steps for the biosynthesis of the melanin-like pigment in which Pks3, Pks4, Pks5 and Cyp4 play an earlier role in the pathway, followed by Vbs1 and Deh1. The fact that the extracts of the deletion mutants of *orf1, orf4, orf5, aox1, omt1* and *pmo1* showed similarity to the P*crg*::*mtf1* strain, illustrates that the proteins encoded by these genes may play a minor role in the biosynthesis of the melanin-like pigment.

**Fig. 2.**
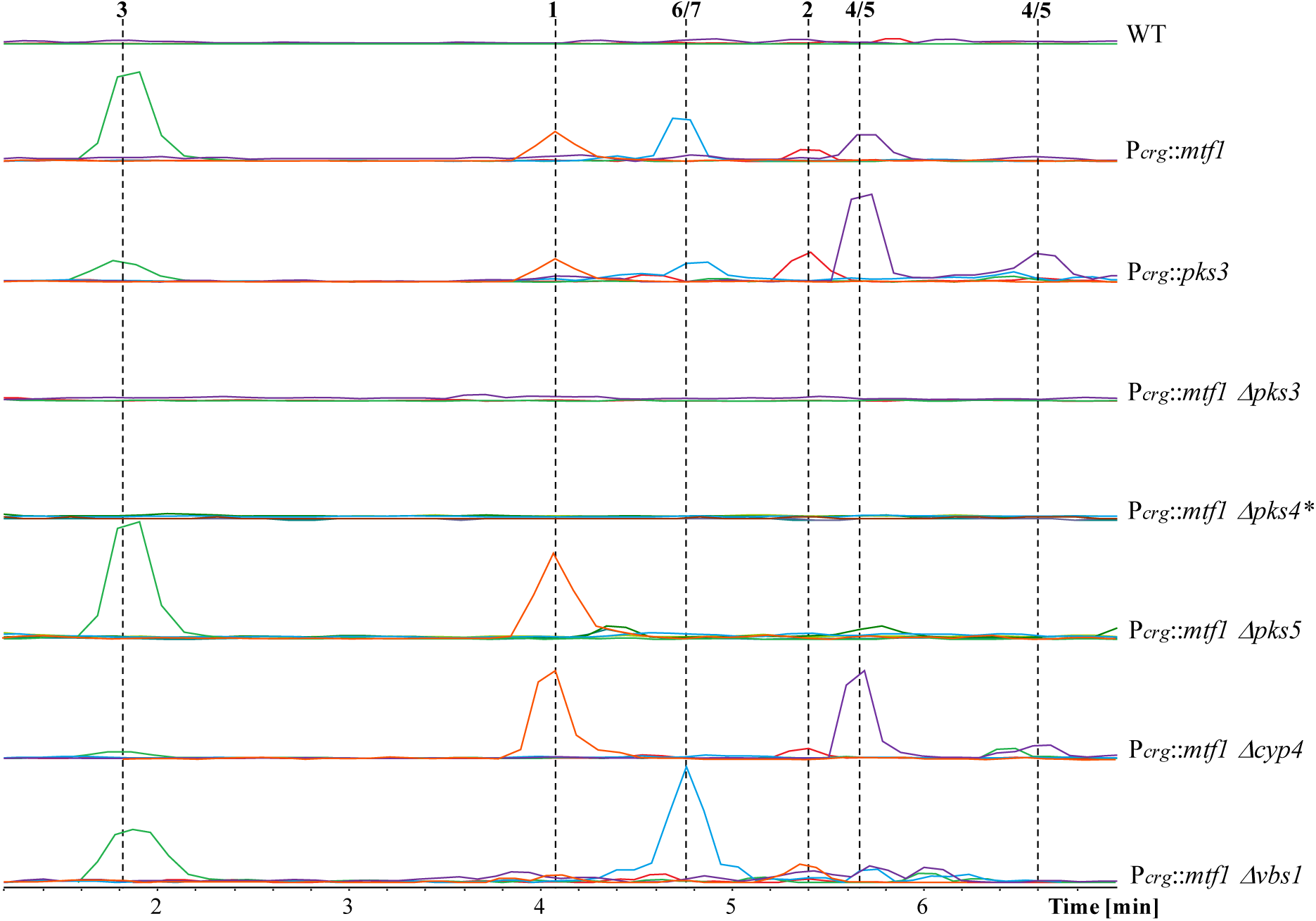
Metabolic profiling of the melanin cluster deletion mutants. Metabolic profiles of the cultured cells from the wild-type, overexpressing strain of *pks3* and deletion mutants of *pks3, pks4, pks5, cyp4* and *vbs1* in the P*crg*::*mtf1* strain. Extracted ion chromatograms (EICs) for all the identified natural products are shown inTable S6. For structures corresponding to the numbered natural products, see Fig. 3B.

### Pks3 and Pks4 are required for the biosynthesis of orsellinic acid (OA)

From our previous results, we observed that deletion of either *pks3* or *pks4* in the P*crg*::*mtf1* strain showed a colorless phenotype and none of the mutants were capable of producing detectable compounds. Additionally, the extracts of P*crg*::*mtf1* Δ*pks5* accumulated OA and TAL but no melanin suggesting the involvement of Pks5 in the processing of OA and or TAL to melanin. Therefore, we asked ourselves whether Pks3 and Pks4 could synthesize together OA and TAL. To achieve this goal, we generated single and double over-expressing strains of *pks3* and *pks4* in the background of the wild-type strain MB215. Each PKS-encoding gene was placed under the control of the arabinose-inducible promoter *crg*, thus generating the following strains: P*crg*::*pks3*, P*crg*::*pks4* and P*crg*::*pks3*+P*crg*::*pks4*. Cultures of the double over-expressing strains displayed similar phenotypes as those observed for their single over-expressed genes. While the chromatograms of the single over-expressing strains P*crg*::*pks3* and P*crg*::*pks4* did not exhibit the presence of compounds, prolonged induction of the strain P*crg*::*pks3*+Pcrg::*pks4* triggered the production of OA (**1**) **(Fig. S3A)**. These findings indicate that only the combined expression of Pks3 and Pks4 are sufficient for synthesizing OA and none of the single PKSs possesses the potential for producing this compound **(Fig. S3A)**. To corroborate this result, we performed an *in vitro* experiment where the strains P*crg*::*mtf1* Δ*pks3* and P*crg*::*mtf1* Δ*pks4*^*\**^ were fed with OA with the primary goal of observing a chemical complementation of the mutants by the addition of this compound due to the bypass of the first biosynthetic reaction in the pathway. As controls, we also included the strains P*crg*::*mtf1* Δ*pks5* and P*crg*::*mtf1* Δ*cyp4*, since according to our previous observations, Pks5 and Cyp4 play a role in downstream reactions to those in which Pks3 and Pks4 are involved. As anticipated, the addition of OA to either P*crg*::*mtf1* Δ*pks3* or P*crg*::*mtf1* Δ*pks4*^*\**^ rescued the phenotype of these mutants to a certain extent compared to the P*crg*::*mtf1* strain. In contrast, the presence of OA did not darken the cultures of the P*crg*::*mtf1* Δ*pks5* and P*crg*::*mtf1* Δ*cyp4* strains **(Fig. S3B)**. The present findings allow us to understand that Pks3 and Pks4 are absolutely required for the synthesis of OA, a compound that is further modified to produce melanin.

## DISCUSSION

Melanin is considered as a multifunctional pigment found in all biological kingdoms (6). In *U. maydis*, melanin plays an important role in survival in harsh environments and pathogenicity during maize-plant infection. Although previous reports in *U. maydis* suggest that two PKS-encoding genes (*pks1* and *pks2*) and a putative laccase (*lac1*) are required for producing pigmented teliospores in maize tumors (16), the mechanism of its biosynthesis in this fungus remains unclear. In this work, we were able to identify a large PKS gene cluster involved in the biosynthesis of melanin in *U. maydis* via an unconventional pathway. Accumulation of melanin was observed upon prolonged induction of the transcription factor *mtf1*, whose activation led to simultaneous up-regulation of 12 genes including three PKSs (*pks3, pks4* and *pks5*), a cytochrome p450 (*cyp4*) and a versicolorin B synthase-like gene (*vbs1*), among others (**Fig. 1**). Deletion approaches of the cluster genes in the P*crg*::*mtf1* strain allowed us to elucidate the sequential steps of the biosynthesis of this pigment. We propose that the initial reactions are catalyzed by Pks3, Pks4 and Pks5 since mutants defective in any of these genes displayed an albino phenotype (**Fig. 1C**). Also, the analysis of the extracts of *pks3* or *pks4* mutants by LC-MS revealed that neither deletion mutant was able to generate any UV-detectable compound, while *pks5* mutants caused the production of two compounds, OA and TAL **(Fig. 2)**. These results suggest that Pks3 and Pks4 act at an early stage of the metabolic pathway followed by Pks5, catalyzing the next downstream reaction. Our observations were further supported by the fact that only co-expression of *pks3* and *pks4* gave rise to the synthesis of OA, a compound that was sufficient to chemically complement the phenotype of the strains P*crg*::*mtf1* Δ*pks3* and P*crg*::*mtf1* Δ*pks4\** in a feeding experiment **(Fig. S3)**. To the best of our knowledge, these findings constitute the first report of a fungus capable of synthesizing OA by the combined function of two PKSs which are also required, together with a third one, in a pathway that leads to the production of melanin. By analyzing the domain structure of these PKSs, it can be noticed that neither Pks3 nor Pks4 possesses the three minimal domains required for being considered as functional PKSs **(Fig. 3A)**, the β-ketoacyl synthase (KS), the acyltransferase (AT) and the acyl carrier protein (ACP) domains (22). Although Pks3 has a similar domain structure when compared to other OA-PKSs (23), the lack of an AT domain in this protein lets us speculate that this activity is probably recruited *in trans* from Pks4, in a similar fashion as described for bacterial *trans*-AT PKS systems (24). In fact such a split-PKS system would be similar to fungal fatty acid synthases (FAS) which are multimeric complexes build from two different FAS enzymes with complementary domain functionality (25). Nevertheless, further experimental evidence has to be provided to understand the reaction mechanism of Pks3 and Pks4 in more detail. Along with OA, TAL was identified in the extracts of the *pks5* mutants, which denotes that this compound is also produced at earlier stages in the same pathway. In *Penicillium patulum*, TAL is synthesized by a 6-methylsalicylic acid synthase in the absence of the reducing agent NADPH, a reaction that requires the condensation of two malonyl-CoA molecules with one acetyl-CoA starter unit, followed by cyclization and aromatization (26, 27). In our model, we are proposing that TAL is synthesized similarly as in *P. patulum* but instead of requiring a single enzyme, this product is afforded by two PKSs just as in the case of OA **(Fig. 3B)**. Biosynthesis of OA and TAL has been previously documented in *Aspergillus terreus*, where the heterologous expression of a single *pks*-gene (*terA*) in *Aspergillus niger* yielded a mixture of these compounds together with 6,7-dihydroxymellein (28).

**Fig. 3.**
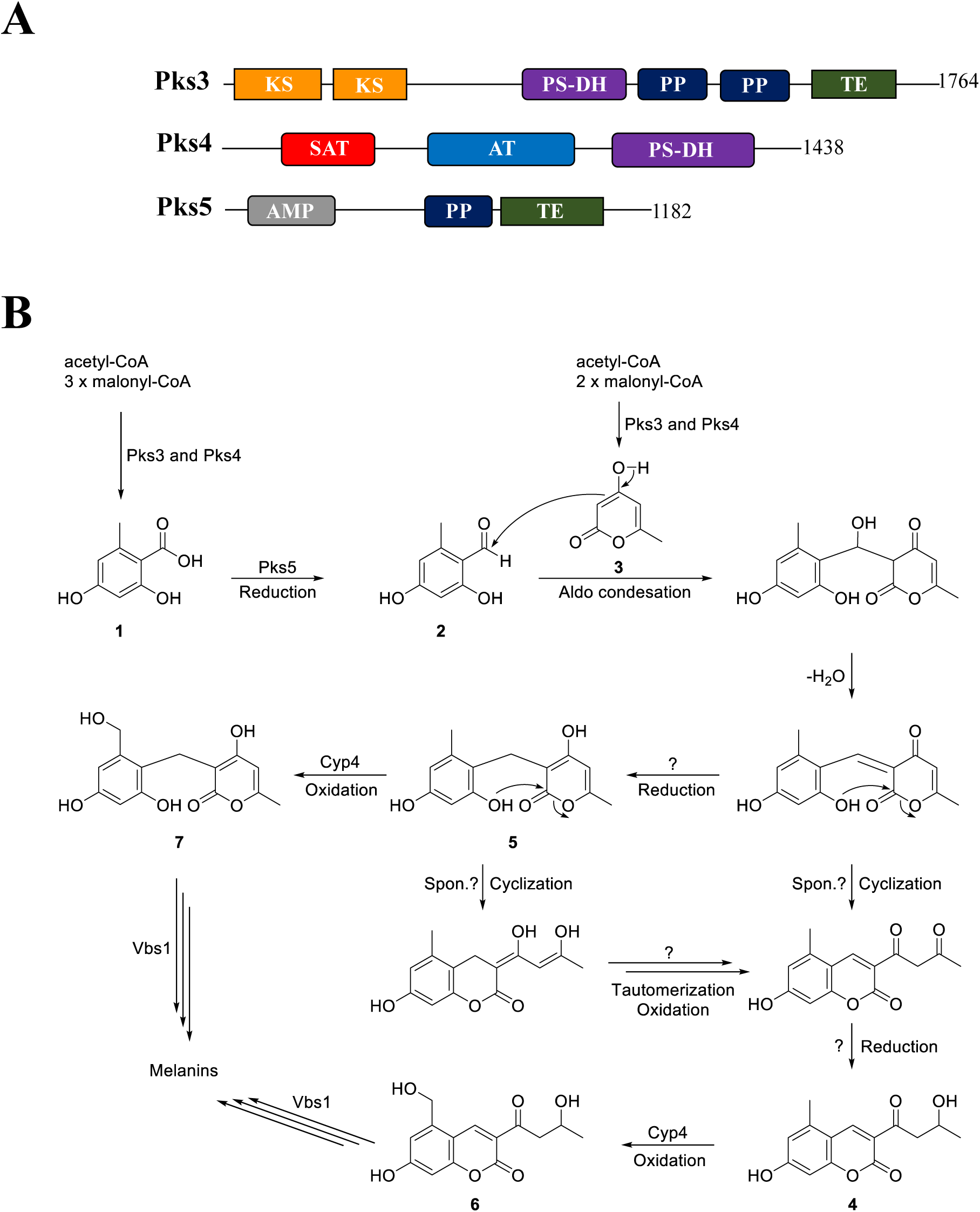
Mechanism of biosynthesis of melanin in *U. maydis*. **(A)** Domain organization of polyketide synthases of the melanin-like gene cluster in *U. maydis*. Protein enzymatic domains are as follows: AT, acyltransferase; AMP, adenylation; KS, β-ketoacyl synthase; NAD, nicotinamide adenine dinucleotide binding; PP, phosphopantetheine attachment site; PS-DH, polyketide synthase dehydratase; SAT, starter unit and TE, thioesterase. Domains were predicted by Protein Families (PFAM) and National Center for Biotechnology Information (NCBI) domain search. Numbers located on the right side of each PKS indicate their length in amino acids, and numbers below each domain represent their localization in the protein. **(B)** Proposed biosynthetic metabolic pathway in *U. maydis*.

On the other hand, the production of orsellinic aldehyde (**2**) is achieved by the reduction of OA, a reaction catalyzed by PKS5. Although the domain structure of PKS5 does not correspond to a canonical PKS either (**Fig. 3A**), its thioesterase domain (TE) appears to be responsible for this reductive step (29).

Together with OA, TAL and orsellinic aldehyde, the coumarin- and 2H-pyrane-2-one-like compounds (**4** and **5**, respectively) were isolated from the *cyp4* disrupted mutant (**Fig.2**), which showed a slightly darker phenotype if compared to any of the *pks*-deletion mutants (**Fig. 1C)**. Such compounds have been reported to have antimicrobial properties in other fungi (30, 31), which makes us think that they could have a similar role in *U. maydis*. Since cytochrome P450s act as terminal monooxygenases in a range of biochemical reactions (32), we consider that compounds **6** and **7** are synthesized upon Cyp4 dependent oxidation of **4** and **5**, respectively **(Fig. 3B)**.

Disruption of *vbs1* caused a yellowish phenotype **(Fig. 1C)** and accumulation of compounds **6** and **7**, the proposed products of the reactions catalyzed by Cyp4 **(Fig. 3).** Although we found no reports of involvement of versicolorin B synthases in the biosynthesis of melanin, deletion of a versicolorin B synthase-encoding gene that is part of the aflatoxin pathway of *A. parasiticus*, resulted in the accumulation of 5’-oxoaverantin and yielded a yellowish pigment (33).

On the other hand, none of the deletion mutants of the co-regulated cluster genes *omt1, pmo1, deh1, orf1, orf4* and *orf5* gave a colorless phenotype **(Fig. 2)**, indicating that these genes might encode for proteins that act at the end of the metabolic pathway as tailoring enzymes of the actual melanin. No conserved domains were detected in *orf1, orf4* and *orf5*, neither similarities with proteins from other fungi. However, a closer analysis of their amino acid sequences shows a certain degree of similarity among them, which makes us suspect that these genes are paralogous genes residing within the same cluster.

In fungi, biosynthetic gene clusters are often linked to more than one compound, but only in few cases, the same kind of compound is linked with two distant and unrelated clusters from the same genome. One example is *Botrytis cinerea*, whose melanogenesis process is based on two developmentally regulated PKS-encoding genes (BcPKS13 and BcPKS12) that are capable of generating the same intermediate T4HN in conidia and sclerotia, respectively. Furthermore, this compound is converted to DHN by the same set of enzymes, resulting in the production of melanin (34). The work presented here together with the data shown by García-Pedrajas and collaborators (16), suggest that *U. maydis* harbors two gene clusters that are involved in the synthesis of melanin. However, no enough evidence suggests that the final product corresponds to the same kind of melanin. RNA-seq data of the FB1 strain showed that the expression levels of *pks3, pks4* or *pks5* are low in axenic culture as well as during maize plant infection (personal communication, Daniel Lanver), which strengthens the evidence that *pks1* and *pks2* are the major players in melanization during maize plant infection. In a previous study, Jonkers and collaborators determined the metabolic and transcriptomic changes in *U. maydis* and *Fusarium verticillioides* when both fungi were co-cultivated. Seven of the melanin cluster genes (*pks5, orf1, pks4, vbs1, omt1, pmo1* and *cyp4*) showed slightly higher expression levels during co-cultivation with *F. verticillioides* compared to the single cultures, suggesting that *U. maydis* could activate those genes in response to the presence of competitors (35). Although our studies reveal a novel mechanism for the synthesis of melanin in fungi, further studies need to be performed in order to shed light on the biological function of this cluster in *U. maydis*, as well as the complexity of its unique PKS system.

## MATERIALS AND METHODS

### Strains and Cultivation Conditions

*Escherichia coli* strain TOP10 (Invitrogen) was used for cloning purposes and cultivated in dYT [1% (w/v) yeast extract, 1.6% (w/v) tryptone, 0.5% (w/v) NaCl] liquid medium at 37 °C with shaking or on dYT-agar plates (13 g/l). Ampicillin was used at a concentration of 100 μg/ml. *Ustilago maydis* strains used in this study are listed in **Table S4**, they are derivates of the haploid strains MB215 (*a2 b13*) or FB1 (*a1 b1*). Cells were grown at 28 °C in liquid yeast extract-peptone-sucrose (YEPS_light_, 1% yeast extract, 2% peptone, 2% sucrose) or on solid potato dextrose (PD) agar containing 1.5% (w/v) of Bacto agar. For the selection of transformants, PD plates containing the appropriate antibiotic were used. For the induction of the P*crg* promoter, strains were grown at 28°C in 30 ml of Yeast Nitrogen Base (YNB) liquid medium, pH 5.8, containing 0.1% ammonium sulfate and 5% glucose, to exponential phase. Cells were collected by centrifugation, washed twice with ddH_2_O and resuspended in fresh medium with 0.1% ammonium sulfate and 5% arabinose as a sole carbon source. Cultures were grown with constant shaking for an additional 4 h (RNA extraction) or 96 h (preparation of the extracts).

### Standard Molecular Methods

Standard molecular biology methods were used as previously described (36). Transformation of *U. maydis* followed the protocol of Schulz and collaborators (37). Transformation of *Saccharomyces cerevisiae* was done according to Gietz and Woods (38). *U. maydis* chromosomal DNA was isolated as described (39). RNA was isolated from cells grown in liquid medium using TRIzol Reagent (Life Technologies, Darmstadt, Germany) as described by the manufacturer. For Southern blot analysis, genomic DNA was digested with appropriate restriction enzymes (New England Biolabs and Fermentas), separated on 1% (w/v) agarose gels and transferred to Hybond-N^+^. For Northern blot analysis, 20 μg of total RNA was loaded per lane. Hybond-N^+^ membranes were stained with Methylene Blue (200 mg/l in 300 mM Na-acetate, pH 5.4-5.6) to detect rRNA as a loading control. For radioactive labeling of DNA, the megaprime DNA labeling kit (Amersham Biosciences, Braunschweig, Germany) was used. Specific α-^32^P-dCTP labeled probes for Southern and Northern blots were prepared by PCR amplification with their respective primer pair as indicated in **Table S1**. PCR reactions were performed using the DNA polymerase Phusion (lab preparation) for short fragments (< 5 kb) or KOD Xtreme Hot Start polymerase (Novagen) for longer fragments (>5 kb). All PCR products were cleaned up (Geneaid, Taipeh, Taiwan) before digestion. Ligation procedures were carried out with T4-DNA-ligase with supplemented buffer (Roche, Mannheim, Germany).

### Genetic Manipulation of *U. maydis* and Transformant Analysis

Deletion constructs were generated by using the yeast drag and drop method (40). The 5’- and 3’-non-coding regions of the candidate genes were amplified using the primer combinations LBfw/LBrv and RBfw/RBrv listed in **Table S1**. The entire ORFs of the genes were replaced by a hygromycin- or geneticin-cassette except for *pks4, pks5* and *orf1*, where only 0.4-0.5 kb of each gene was deleted. For *pks5* (*UMAG_04095*), 1 kb downstream of the 5’-non-coding region was used as a left border and amplified with the primer pair MI287_pks5_LB_fw / MI288_pks5_LB_rv, while the region located at 1.5-2.5 kb downstream of the start codon was used as a right border (MI289_pks5_RB_fw / MI290_pks5_RB_rv. In the case of *pks4* (*UMAG_04097*), the downstream region of the stop codon spanning from 2-3 kb was taken as a left border (MI469_pks4_LB_fw / MI470_pks4_LB_rv), whereas the right border included 0.77 kb upstream and 0.22 kb downstream of the 3’-non-coding region (MI471_pks4_RB_fw / MI472_pks4_RB_rv). Deletion construct of *orf1* was assembled by amplifying the left and right borders with the primer pair MI938_orf1_LB_fw / MI939_orf1_LB_rv and MI940_orf1_RB_fw / MI941_orf1_RB_rv, respectively. Left border fragment comprised 0.78 kb of the 5’-non-coding region and the first 0.25 kb of the ORF of *orf1*, while the right border was constituted of the last 0.1 kb of the ORF and 0.88 kb of the 3’-non-coding region. Resistance cassettes were SfiI fragments of the plasmids pMF1-h (hygromycin, H) and pMF1-g (geneticin, G) (41). Flanks, hygromycin- or geneticin-resistance cassette and the KpnI/BamHI pRS426 cut vector (42) were assembled using homologous recombination in *S. cerevisiae*. Cloned deletion constructs were verified by sequencing, excised with SspI and used for transformation of P*crg*::*mtf1* protoplasts. In order to generate the *U. maydis* over-expressing strains, plasmids pCRG-Mtf1-Tnos-Cbx, pCRG-Mtf2-Tnos-Cbx, pCRG-Pks3-Tnos-Cbx and pCRG-Pks4-Tnos-Cbx, were constructed by replacing the 0.7 kb XmaI/NotI *egfp* gene fragment of the pCRG-GFP-Ala6-MXN with a XmaI/NotI digested PCR products amplified with the primers: MG700_mtf1_XmaI_fw / MG703_mtf1_NotI (*UMAG_04101*), MG554_mtf2_XmaI_fw /MG551_mtf2_NotI_rv (*UMAG_11110*), MH701_pks3_XmaI_fw / MH702_pks3_NotI_rv (*UMAG_04105*) and MI593_pks4_XmaI_fw /MH704_pks4_NotI_rv (*UMAG_04097*), respectively. The purified PCR product was cloned into the vector pCRG-GFP-Ala6-MXN digested with the same restriction enzymes. For constructing the plasmid pCRG-Pks3-Tnos-Cbx-G418, the P*crg* promoter and the ORF of *pks3* were amplified from the plasmid pCRG-Pks3-Tnos-Cbx with the primer pair: MI985_crg_SbfI_fw / MI986_pks3_AflII_rv, respectively **(Table S1)**. The PCR products were cloned into the pJA2880 plasmid, which had been digested with SbfI and AflII to remove the 1.7 kb pETEF-*egfp* insert. Prior transformation in *U. maydis*, over-expressing plasmids were linearized with SspI and integrated into the *ip* locus by homologous recombination. For the selection of the transformants, PD plates with 200 μg/ml of hygromycin, 200 μg/ml of geneticin or 2 μg/ml of carboxin were used. A successful homologous replacement was verified by Southern blot for all generated strains. In addition, Northern blot analysis was performed to analyze the expression of the desired genes.

### Orsellinic Acid Feeding Experiments

Selected *U. maydis* strains were inoculated in 3 ml of YEPS_light_ medium and incubated at 28 °C ON with constant shaking. Afterward, the cells were diluted to an OD_600_=0.2 in 30 ml of YNB pH 5.8 with 5% glucose and 0.1% ammonium sulfate until an OD_600_=0.6 was reached. Subsequently, the cells were washed twice with dH_2_O and transferred to either 4 ml (feeding with OA) of YNB liquid medium containing 0.1% ammonium sulfate and 5% of glucose (control) or arabinose. Cultures were incubated with shaking for 96 h at 28 °C. For the feeding experiment with OA, the 4 ml cultures were grown in the presence or absence of 0.5 mM of orsellinic acid (Alfa Aesar, Heysham, England) in DMSO.

### Metabolite Extraction Analysis and Structure Elucidation

Pelleted cells of 30 ml liquid cultures were transferred to 50 ml glass beakers and stirred with 20 ml acetone for 1 h. The extracts were filtered into 100 ml round bottom flasks and the pellets were washed with a further 10 ml acetone before being concentrated to dryness in a rotary evaporator. Dry extracts were dissolved in 2 ml methanol and transferred to 4 ml glass vials prior of being concentrated to dryness in a speed vacuum (Speed Vac-Concentrator, Eppendorf, Hamburg) for 2 h at 28 °C. Extracts were dissolved in 1 ml of methanol (HPLC grade) and centrifuged (16,000 rpm, 20 min) to ensure that no particles were injected into the HPLC instrument. Finally, 100 μl of the supernatant was transferred to HPLC plastic vials and subjected to analysis. For large-scale cultures (> 1L), the whole volume was placed into a 2 L flask with 30 g of Amberlite XAD-16 beads (Sigma-Aldrich) agitated at 200 rpm for 2 h at 4°C. Afterward, the amberlite beads were harvested at −20 °C for further analysis. HPLC-MS analysis was performed with a Dionex UltiMate 3000 system coupled to a Bruker AmaZon X mass spectrometer and an Acquity UPLC BEH C18 1.7 μm RP column (Waters).

### Bioinformatic DNA Sequence Analysis

All DNA and protein sequences were retrieved from the National Center for Biotechnology Information (NCBI) database (NCBI; www.ncbi.nlm.nih.gov) and MIPS *Ustilago maydis* Data Base (MUMDB; mips.helmholtz-muenchen.de/genre/proj/ustilago). Homology searches were performed using BLASTp with default settings (http://blast.ncbi.nlm.nih.gov/Blast.cgi).

## ACCESSION NUMBERS

*pks5* (*UMAG_04095)*: XM_011392301; *orf1*(*UMAG_04096*): XM_011392302; *pks4* (*UMAG_04097*): XM_011392303; *orf2* (*UMAG_04098*): XM_011392304; *mtf2* (*UMAG_11110*): XM_011392495; *orf3* (*UMAG_04100*): XM_011392305; *mtf1* (*UMAG_04101*): XM_011392306; *aox1* (*UMAG_11111*): XM_011392496; *vbs1* (*UMAG_11112*): XM_011392497; *orf4* (*UMAG_04104*): XM_011392307; *pks3* (*UMAG_04105*): XM_011392308; *omt1* (*UMAG_04106*): XM_011392309; *pmo1* (*UMAG_04107*): XM_011392310; *orf5* (*UMAG_12253*): XM_011392513; *cyp4* (*UMAG_04109*): XM_011392311; *deh1* (*UMAG_11113*): XM_011392498.

## ACKNOWLEDGMENTS

The authors are grateful to Zakaria Cheikh, Victoria Challinor and Lisa Rosenbecker, who provided insight and expertise that assisted this research. We also thank Marc Strickert for his guidance in the identification of potential *U. maydis* gene clusters.

## FUNDING INFORMATION

Work in the Bölker lab was supported by DFG funded grant SFB987 and by the LOEWE Zentrum SYNMIKRO funded by the state of Hesse.

Work in the Bode lab was supported by the LOEWE Schwerpunkt Mega Syn and the LOEWE Zentrum TBG, both funded by the State of Hesse.

