## Supplementary material for "An Unconventional Melanin Biosynthetic Pathway in *Ustilago maydis*": Fig. S1

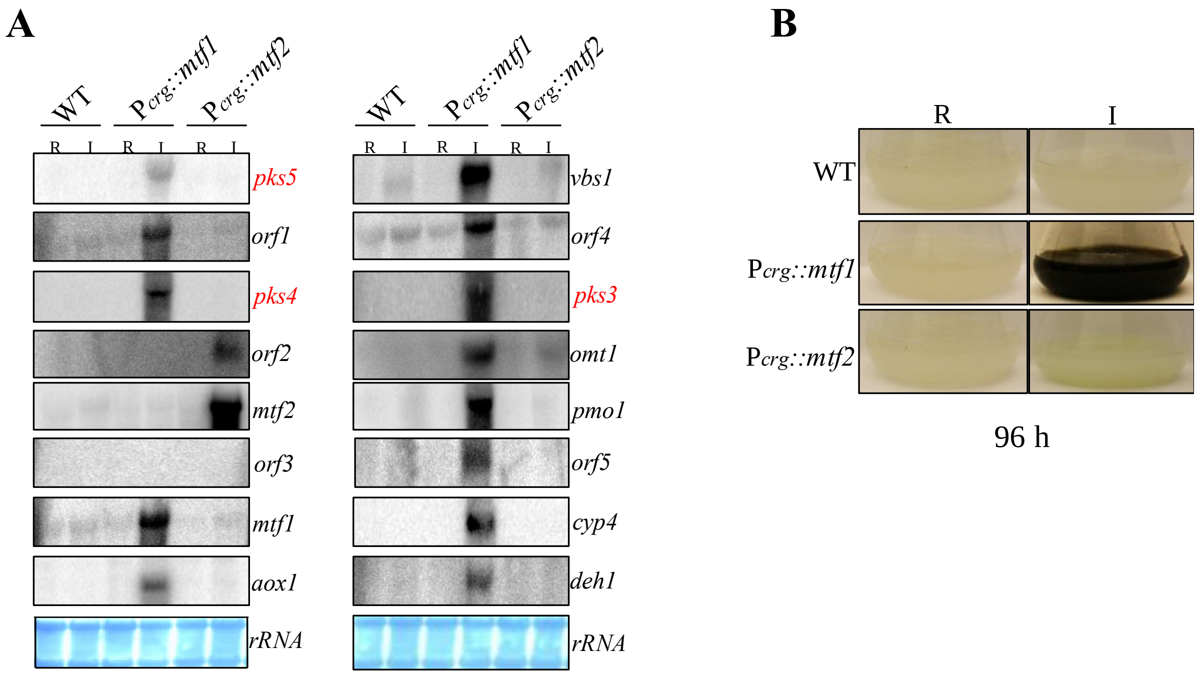


**Fig. S1. Over-expression of the transcription factor *mtf1* stimulates transcription of the melanin cluster genes. (A)** Northern blot analysis of the WT, P*crg*::*mtf1* and P*crg*::*mtf2* strains grown during inducing (I) or repressing (R) conditions for 4 h. Northern blot hybridization probes are indicated on the left side. The bottom panel shows methylene blue-stained rRNA as a control for loading. **(B)** After prolonged induction of *mtf1*, cells produce a dark pigment, while cells grown under repressing conditions remain light.
