## Supplementary material for "An Unconventional Melanin Biosynthetic Pathway in *Ustilago maydis*": Fig. S2

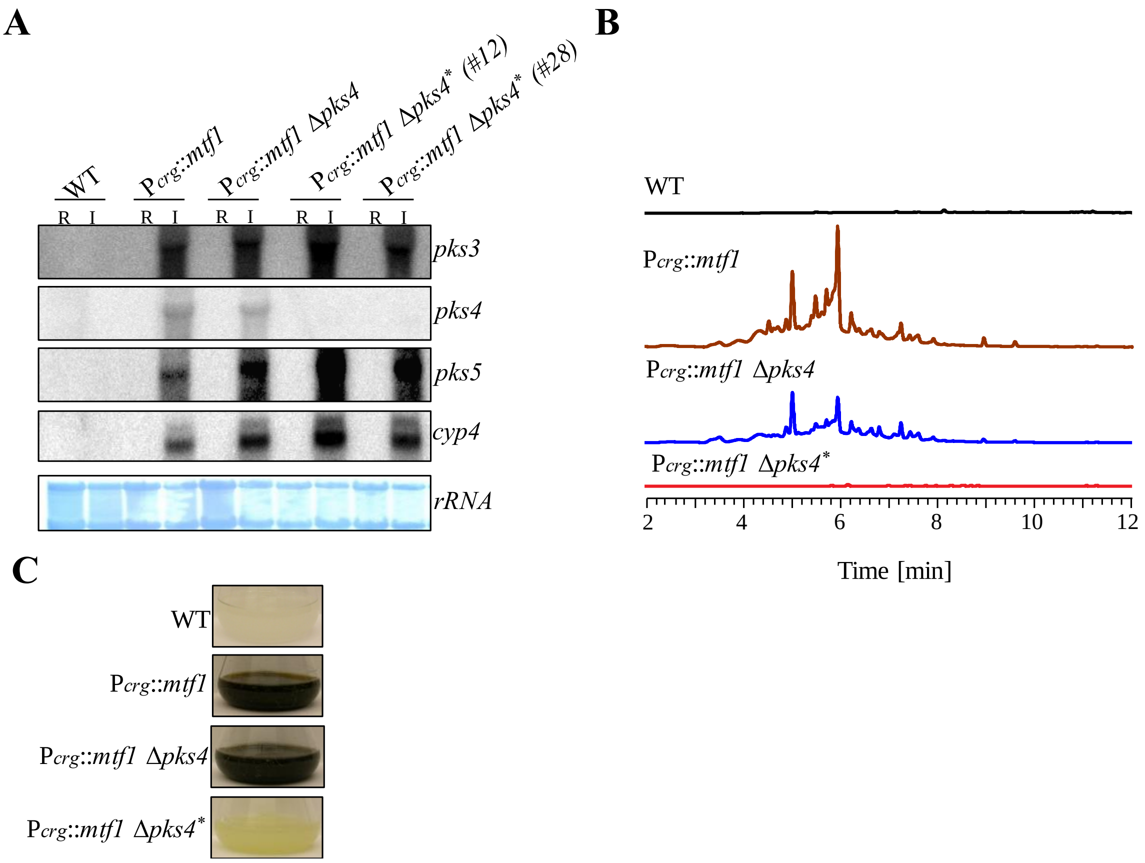


**Figure S2. Single deletion of *pks4* in the P*crg*::*mtf1* strain has no effect in melanin production. (A)** Expression analysis of *pks3*, *pks4*, *pks5* and *cyp4* genes in the induced cultures of the strains P*crg*::*mtf1* Δ*pks4* and P*crg*::*mtf1* Δ*pks4**. All tested strains were grown under repressing (R) and inducing (I) conditions for 4 h at 28 °C. Northern blot was hybridized with the probes indicated on the right side. Lower panel shows methylene blue-stained ribosomal RNA as an indicator of RNA integrity. **(B)** HPLC chromatograms (272 nm) of the wild-type (MB215), P*crg*::*mtf1* Δ*pks4* and P*crg*::*mtf1* Δ*pks4** strains grown in inducing for 96 h at 28 °C in inducing medium. **(C)** Phenotypes of the strains analyzed in B.
