## Supplementary material for "An Unconventional Melanin Biosynthetic Pathway in *Ustilago maydis*": Fig. S3

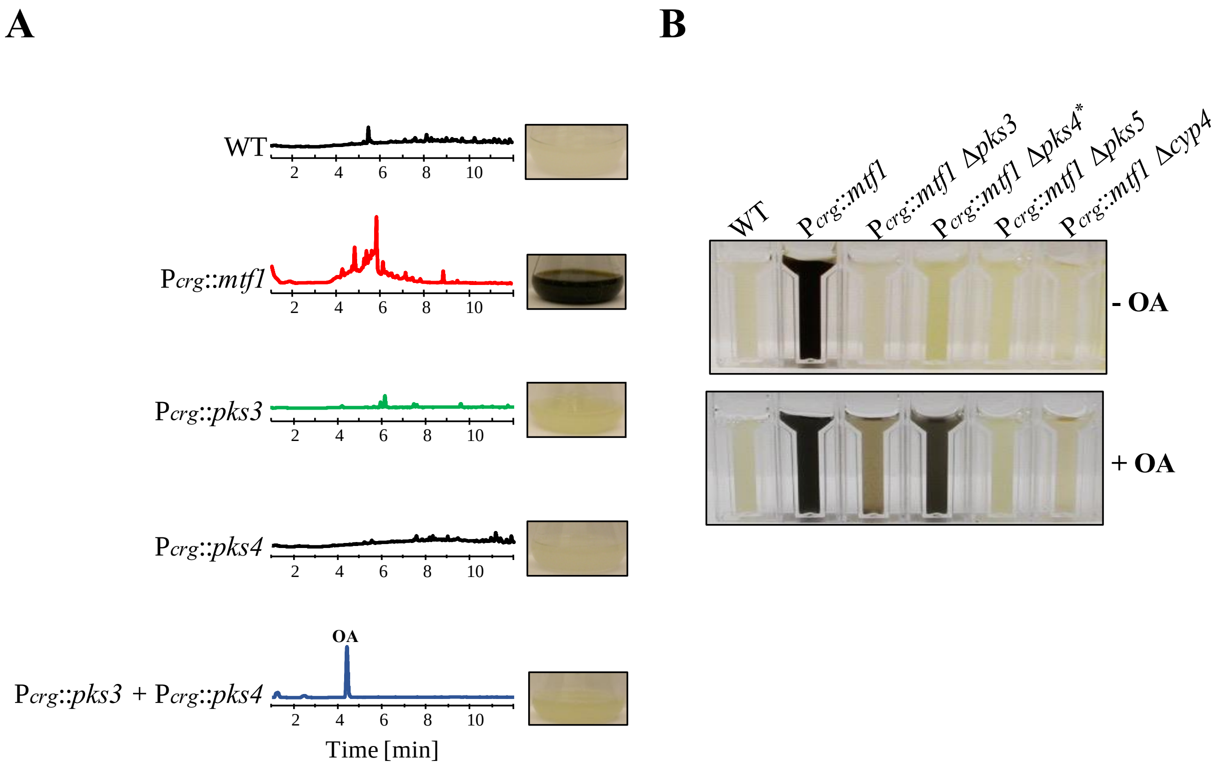


**Fig. S3. Co-expression of *pks3* and *pks4* produces orsellinic acid (OA). (A)** Metabolic profiles of the single (P*crg*::*pks3* and P*crg*::*pks4*) and double (P*crg*::*pks3*+P*crg*::*pks4*) over-expressing strains were analyzed after 96 h of growth in inducing medium at 28 °C. The combined expression of *pks3* and *pks4* produces one single peak identified as OA. **(B)** Feeding of P*crg*::*mtf1* Δ*pks3* and P*crg*::*mtf1* Δ*pks4** with OA can restore the production of the melanin-like pigment. Single deletion mutants P*crg*::*mtf1* Δ*pks3*, P*crg*::*mtf1* Δ*pks4**, P*crg*::*mtf1* Δ*pks5* and P*crg*::*mtf1* Δ*cyp4* were cultivated in inducing medium with the presence (+ OA) or absence (- OA) of 0.5 mM of orsellinic acid for 96 h at 28 °C. Only the feeding of P*crg*::*mtf1* Δ*pks3* and P*crg*::*mtf1* Δ*pks4** with OA could partially complement the melanin-phenotype, whereas P*crg*::*mtf1* Δ*pks5* and P*crg*::*mtf1* Δ*cyp4* were not affected.
