## Supplementary figures and images for "An Unconventional Melanin Biosynthetic Pathway in *Ustilago maydis*"

### Fig. S4

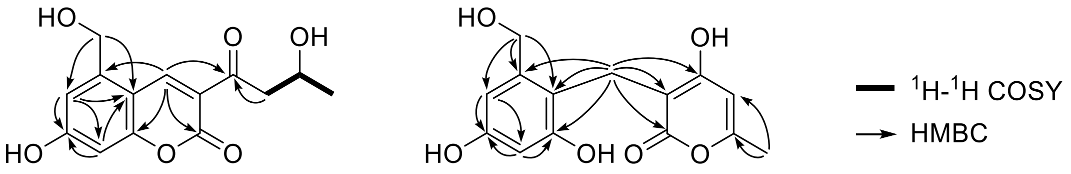


**6** **7**

**Fig. S4. Key 2D NMR correlations of 6 and 7.**

### Fig. S5

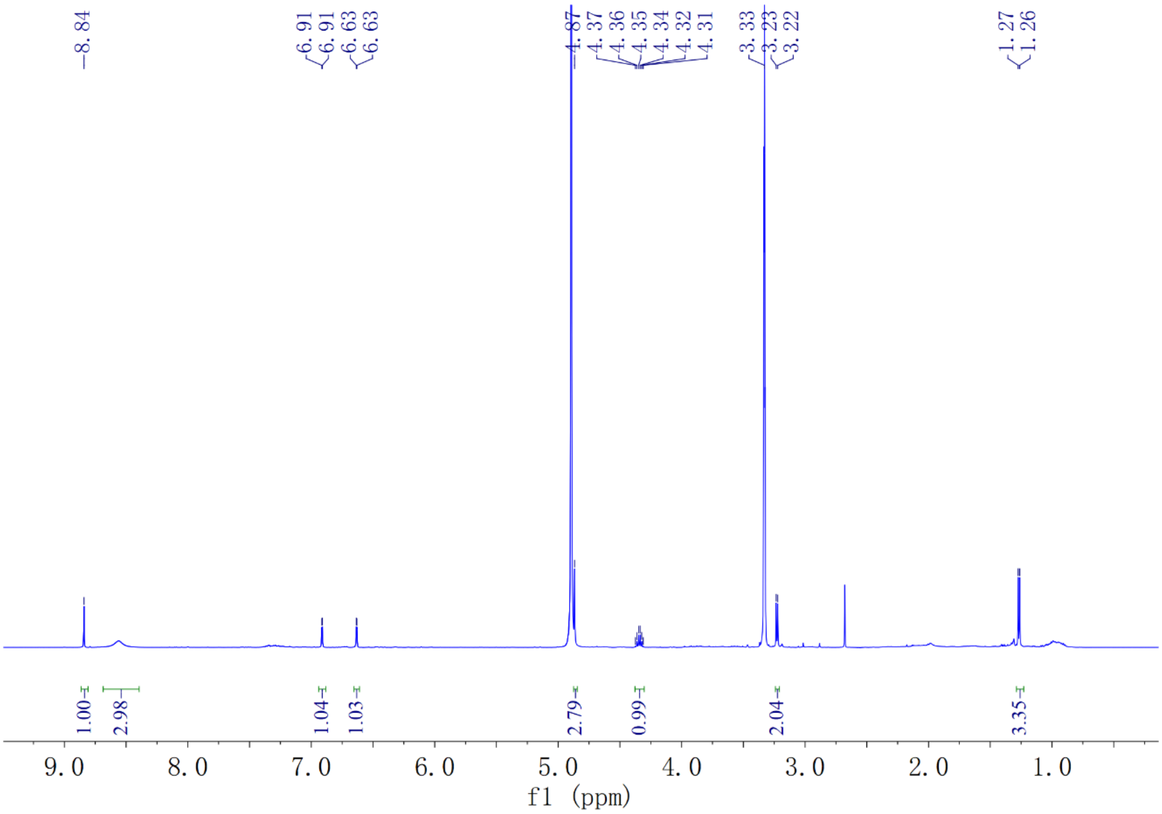


**Fig. S5. [1]H spectrum of 6 in MeOH-*d*$_4$.**

### Fig. S6

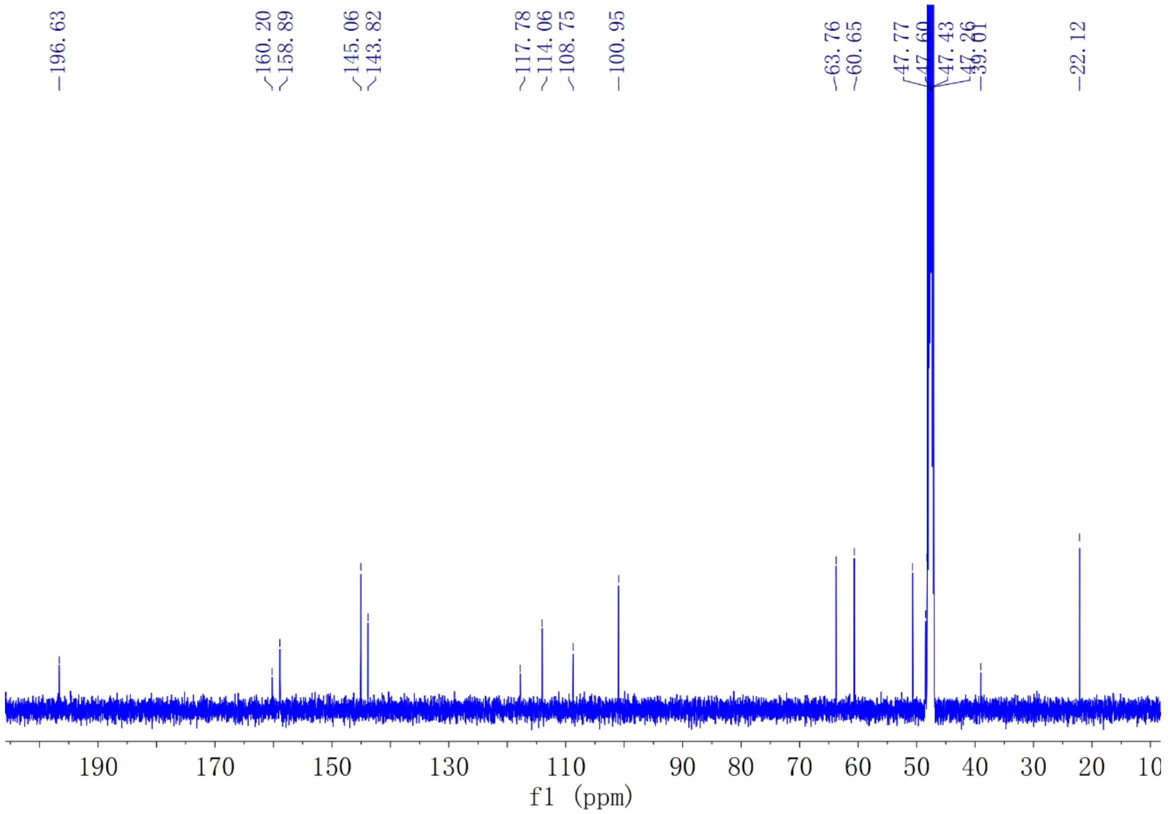


**Fig. S6.** $^{13}$**C spectrum of 6 in MeOH-*d*$_4$.**

### Fig. S7

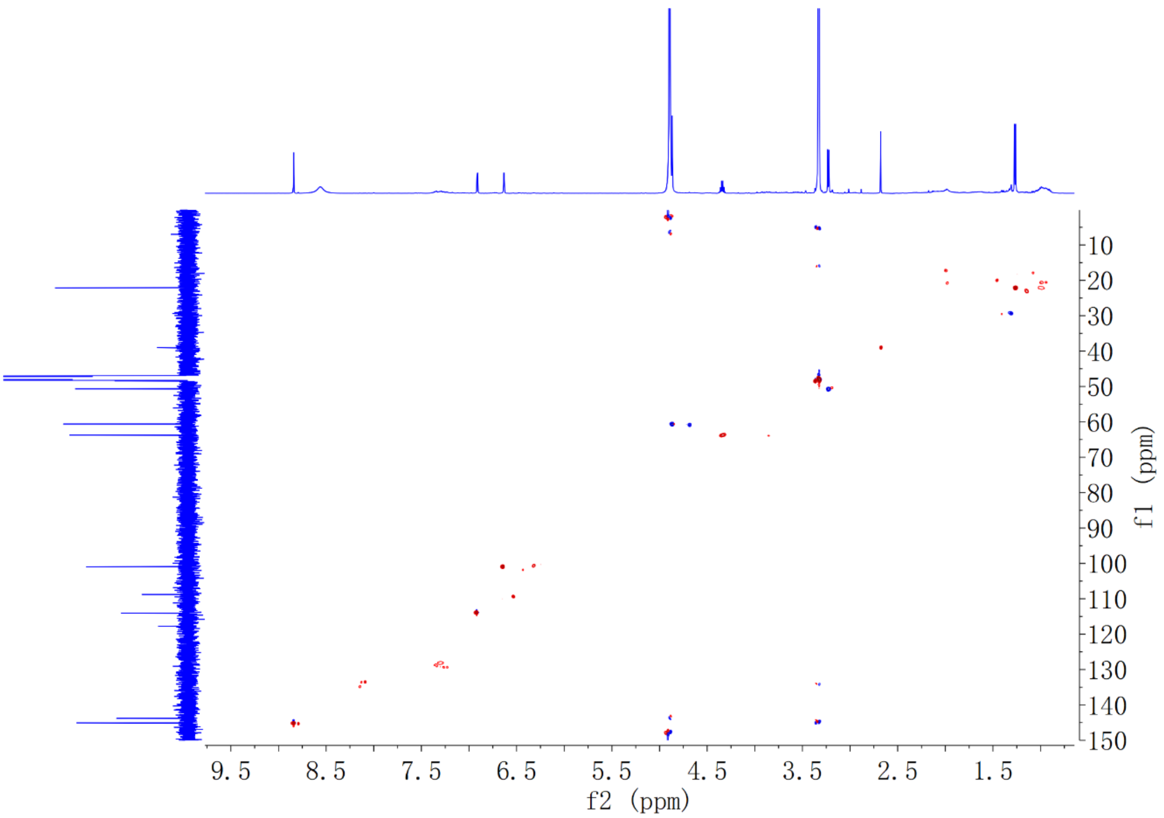


**Fig. S7. HSQC spectrum of 6 in MeOH-*d*$_4$.**

### Fig. S8

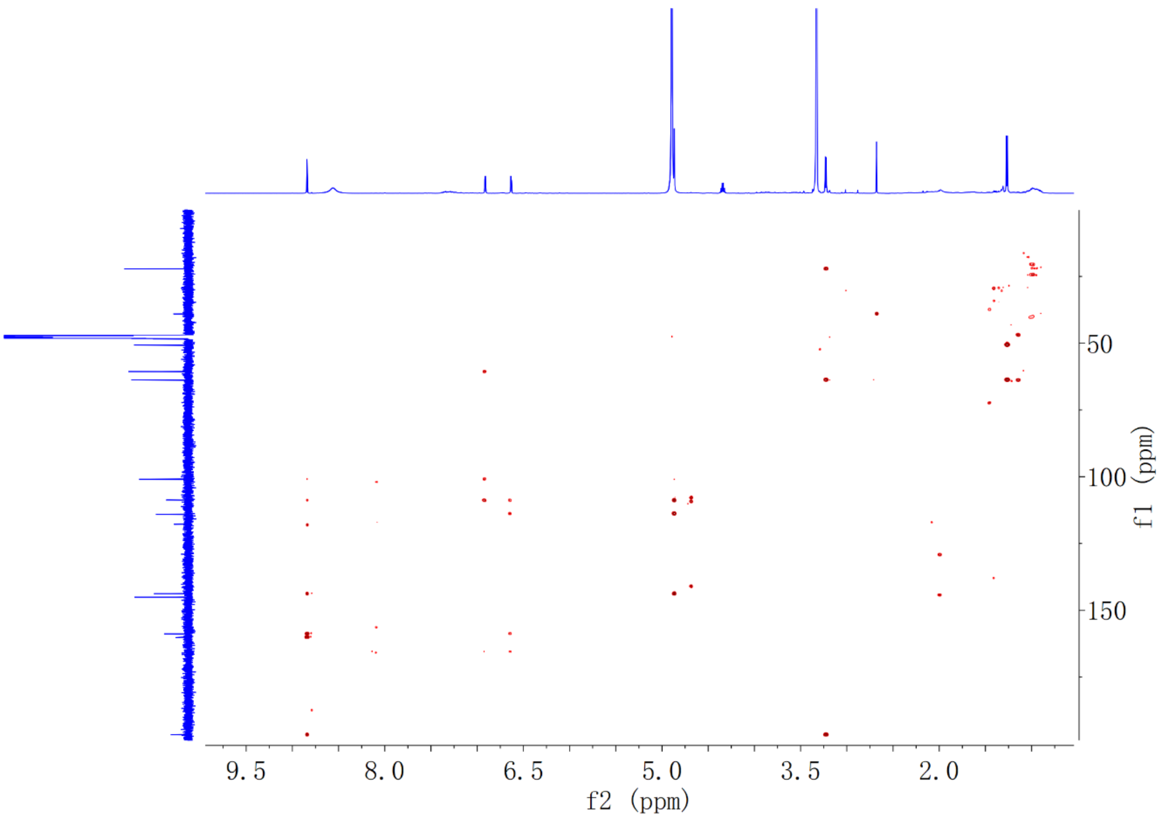


**Fig. S8. HMBC spectrum of 6 in MeOH-$d_4$.**

### Fig. S9

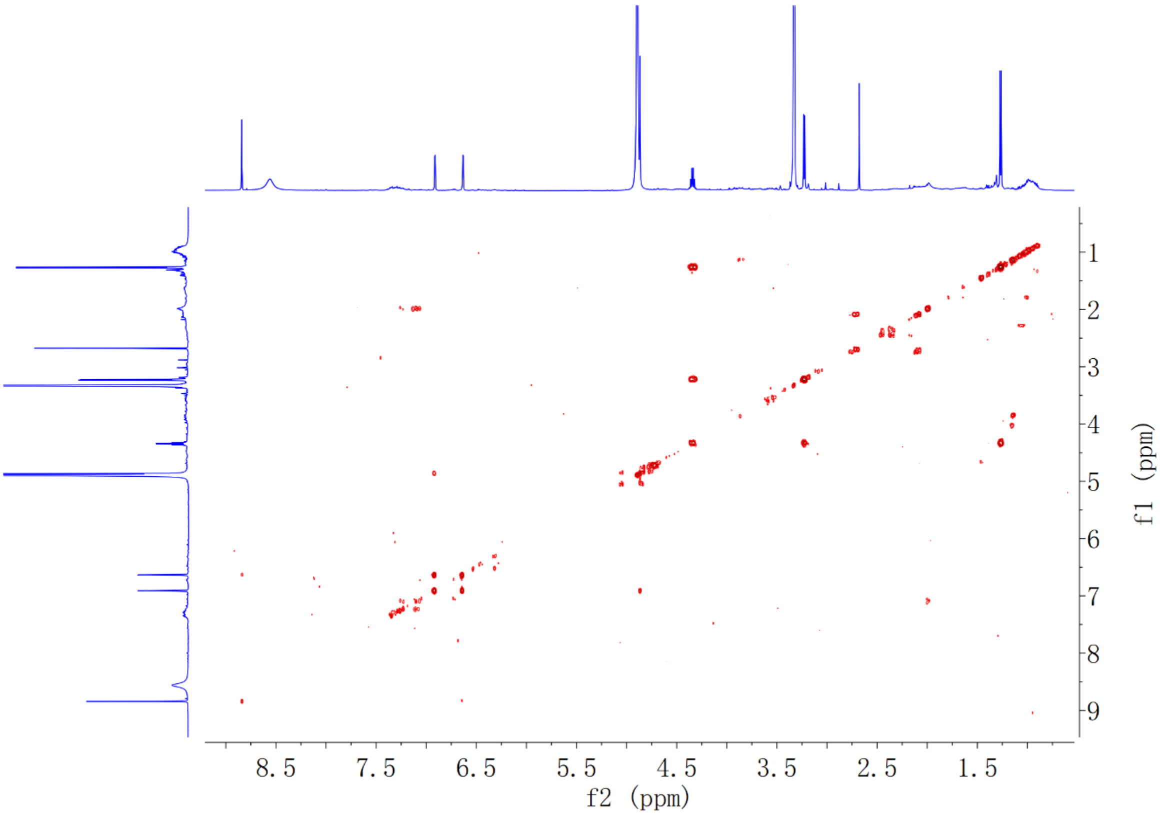


**Fig. S9**. **$^{1}$H-$^{1}$H COSY spectrum of 6 in MeOH-*$d_4$*.**

### Fig. S10

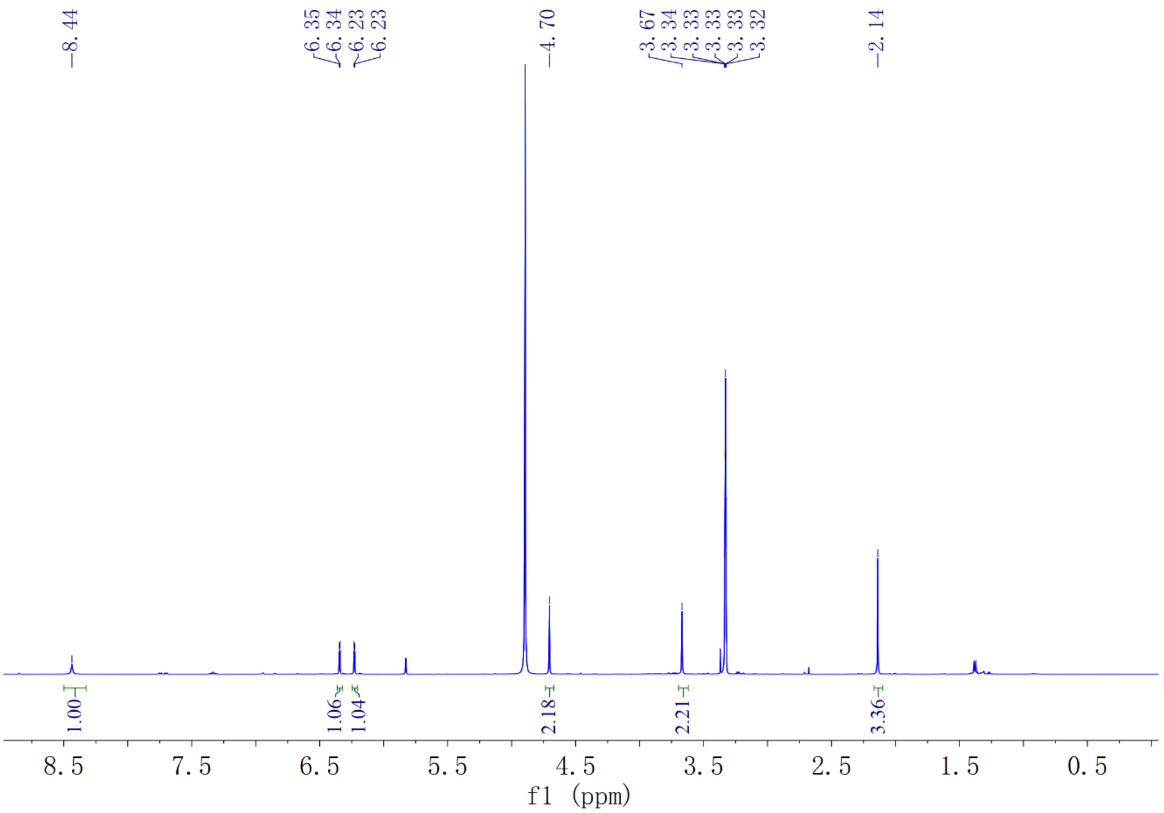


**Fig. S10**. **[1]H spectrum of 7 in MeOH-*d₄*.**

### Fig. S11

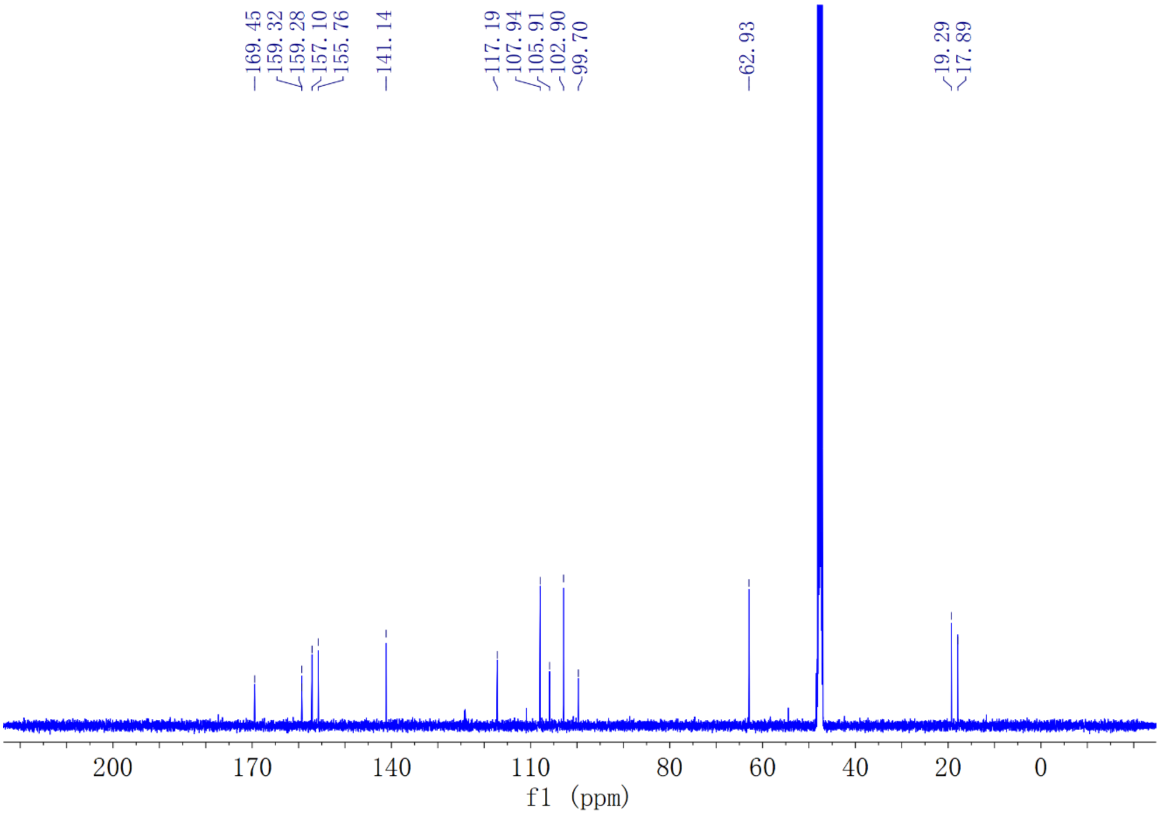


**Fig. S11**. **$^{13}$C spectrum of 7 in MeOH-*d₄*.**

### Fig. S12

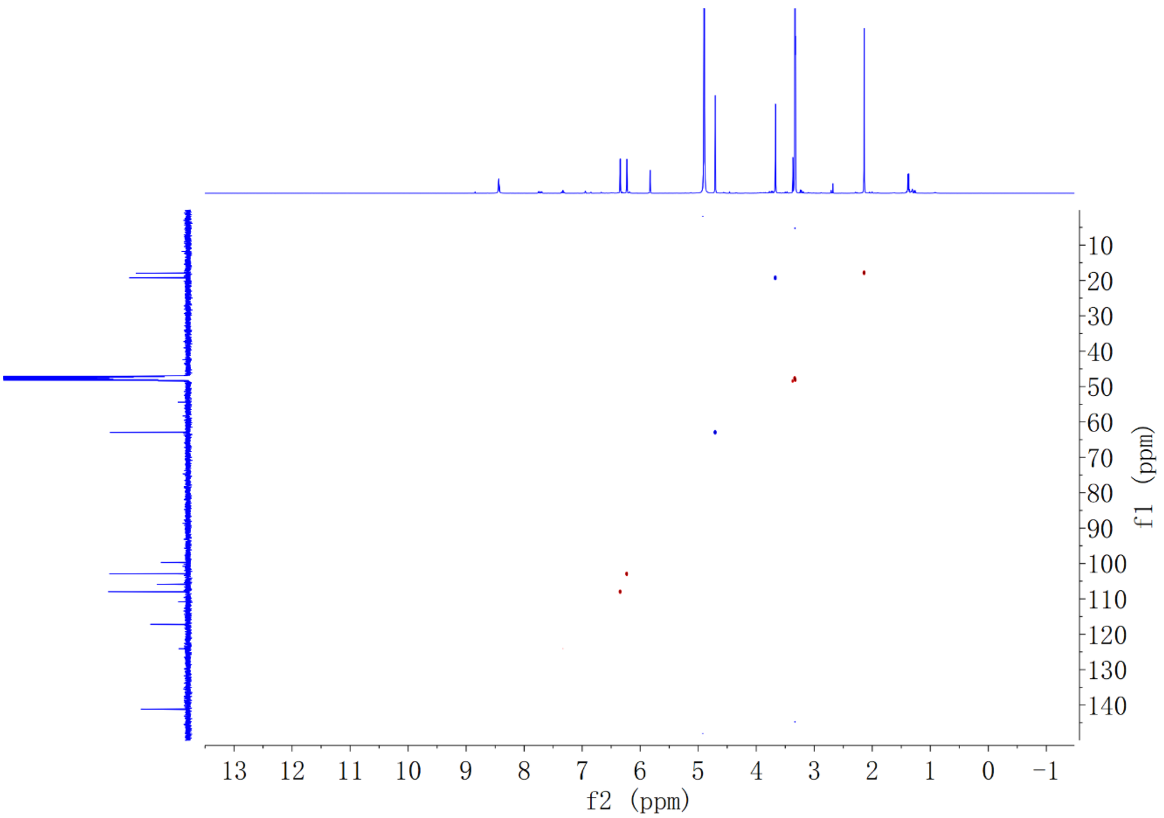


**Fig. S12**. **HSQC spectrum of 7 in MeOH-*d*$_4$.**

### Fig. S13

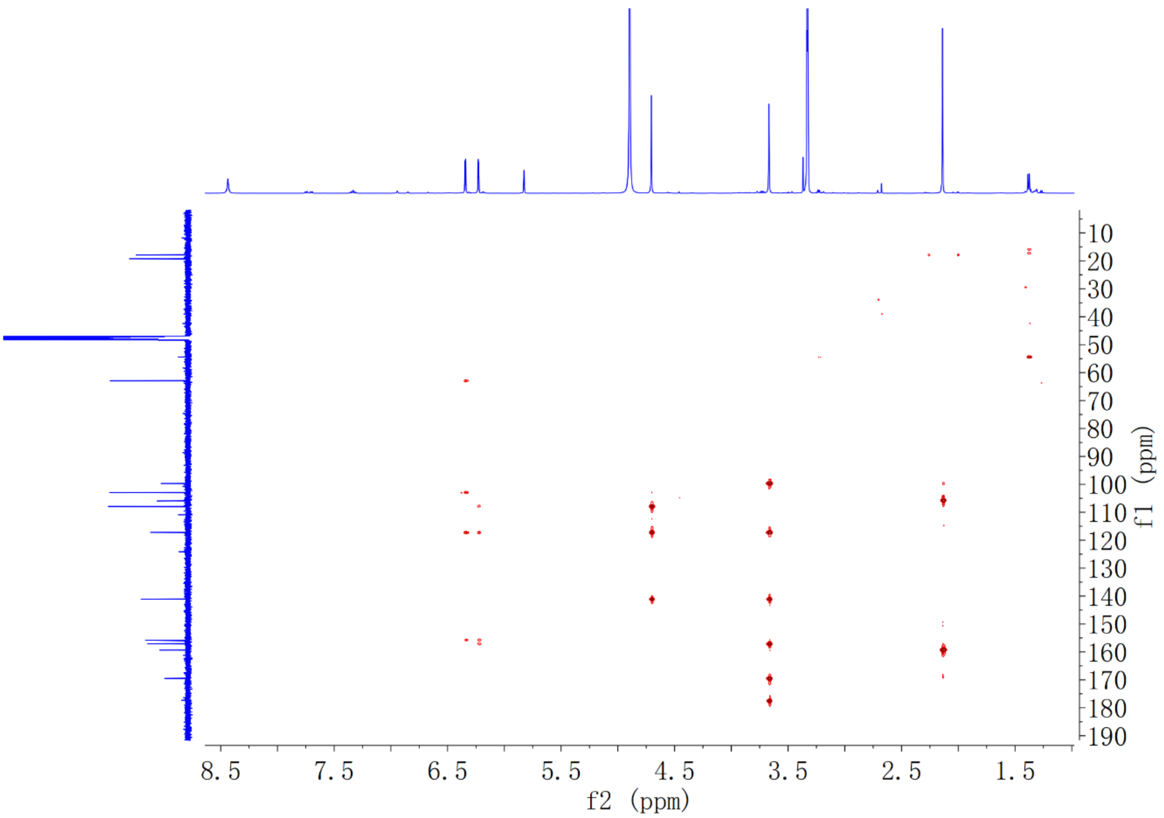


**Fig. S13. HMBC spectrum of 7 in MeOH-*d*$_4$.**

### Fig. S14

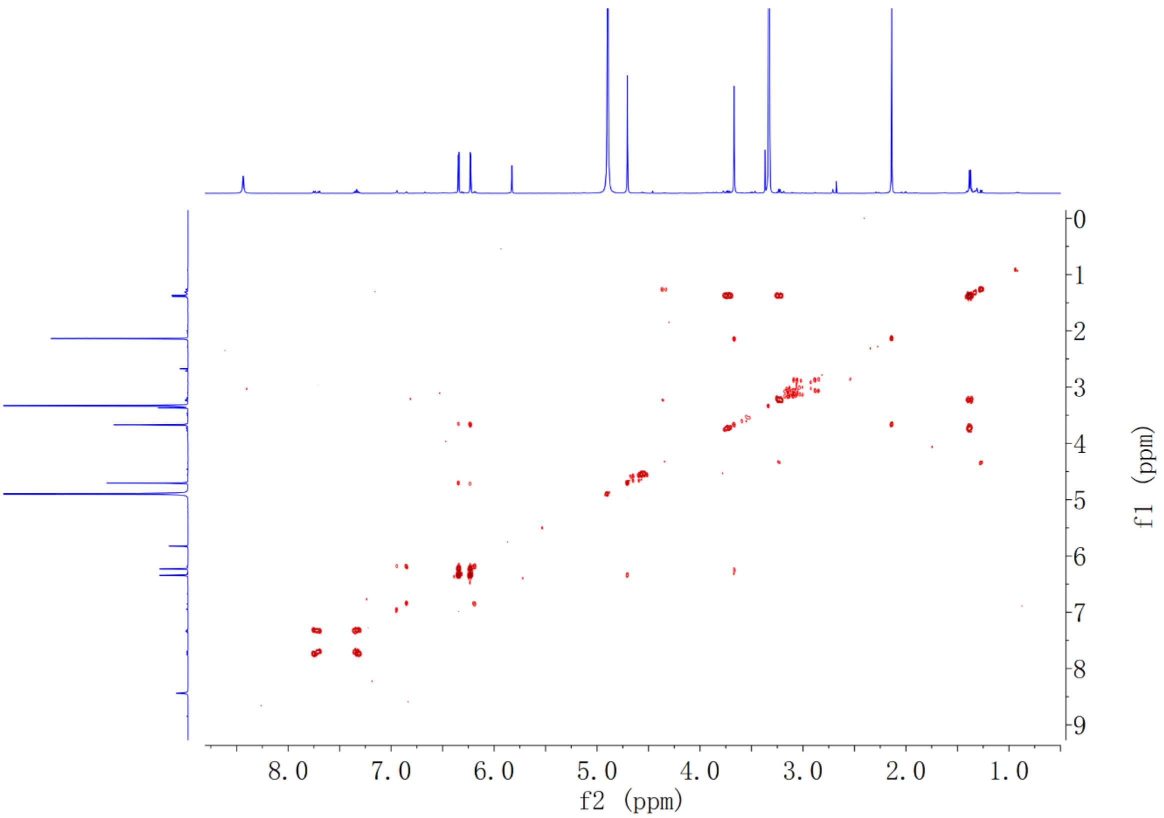


**Fig. S14. $^1$H-$^1$H COSY spectrum of 7 in MeOH-*d*.**
