## Supplementary material for "An Unconventional Melanin Biosynthetic Pathway in *Ustilago maydis*": Table S1

### Supplemental Tables

**Table S1. Oligonucleotides used in this study.**

| # | Name | 5´-->3´ | Template | Function |
|---|---|---|---|---|
| 1 | MI287_pks5_LBfw | gtaacgccagggttttcccagtcacgacgaatatt atgactgtatcacctcctgctccattcg | gDNA of *U. maydis* | flanking regions for deletion |
| 2 | MI288_pks5_LBrv | caattgtcacgccatggtggccatctaggccatc ggcattccgatcgagagcgataggc | gDNA of *U. maydis* | flanking regions for deletion |
| 3 | MI289_pks5_RBfw | gtgcggccgcattaataggcctgagtggccaca atcccaacgctggaggtaatcgggcgcaag | gDNA of *U. maydis* | flanking regions for deletion |
| 4 | MI290_pks5_RBrv | gcggataacaatttcacacaggaaacagcaatat tcgcagatatgtcttaagcaagtgtcgtcc | gDNA of *U. maydis* | flanking regions for deletion |
| 5 | MI938_orf1_LBfw | gtaacgccagggttttcccagtcacgacgaatatt caaagtgaagagagcgtcaagtataagtg | gDNA of *U. maydis* | flanking regions for deletion |
| 6 | MI939_orf1_LBrv | caattgtcacgccatggtggccatctaggccagc ggcatcaatgagggcccttcttcacactc | gDNA of *U. maydis* | flanking regions for deletion |
| 7 | MI940_orf1_RBfw | gtgcggccgcattaataggcctgagtggcccgat tcgttttgtttttcattttcttaccaggatc | gDNA of *U. maydis* | flanking regions for deletion |
| 8 | MI941_orf1_RBrv | gcggataacaatttcacacaggaaacagcaatat tctcccacctcattccattgttggagc | gDNA of *U. maydis* | flanking regions for deletion |
| 9 | MI469_pks4_LBfw | gtaacgccagggttttcccagtcacgacgaatatt gcaggtgtgtttcgcctcgaggacgccatc | gDNA of *U. maydis* | flanking regions for deletion |
| 10 | MI470_pks4_LBrv | caattgtcacgccatggtggccatctaggcccaa ccaccttgtggcccgatactgtctcctggaac | gDNA of *U. maydis* | flanking regions for deletion |
| 11 | MI471_pks4_RBfw | gtgcggccgcattaataggcctgagtggccattc aatctttcactgtggatctacgcgaac | gDNA of *U. maydis* | flanking regions for deletion |
| 12 | MI472_pks4_RBrv | gcggataacaatttcacacaggaaacagcaatat ttctatcttcaacgcatgacgacagtg | gDNA of *U. maydis* | flanking regions for deletion |
| 13 | MG104_mtf2_LBfw | gtaacgccagggttttcccagtcacgacgaatatt cttctatcaacgacttctacaagttgtc | gDNA of *U. maydis* | flanking regions for deletion |
| 14 | MG105_mtf2_LBrv | caattgtcacgccatggtggccatctaggccagc aaagtacaagagcactgcggcattagatc | gDNA of *U. maydis* | flanking regions for deletion |

| | | | | |
|---|---|---|---|---|
| 15 | MG106_mtf2_RBfw | gtgcggccgcattaataggcctgagtggccactt ggctagtcgtgggtcgacgatgccccc | gDNA of *U. maydis* | flanking regions for deletion |
| 16 | MG107_mtf2_RBrv | gcggataacaatttcacacaggaaacagcaatat tcaggacttggcaatgtggatgcgattg | gDNA of *U. maydis* | flanking regions for deletion |
| 17 | MH416_orf3_LBfw | gtaacgccagggttttcccagtcacgacgaatatt gacgaaaccgacacggagcttgatctatatg | gDNA of *U. maydis* | flanking regions for deletion |
| 18 | MH417_orf3_LBrv | caattgtcacgccatggtggccatctaggccgct gtatcatgatgcgttccgaatgaaacataaag | gDNA of *U. maydis* | flanking regions for deletion |
| 19 | MH418_orf3_RBfw | gtgcggccgcattaataggcctgagtggccacc aacagttcccagttaagttactgtacatg | gDNA of *U. maydis* | flanking regions for deletion |
| 20 | MH419_orf3_RBrv | gcggataacaatttcacacaggaaacagcaatat tggccgccaacgcccttttcactagac | gDNA of *U. maydis* | flanking regions for deletion |
| 21 | MG108_mtf1_LBfw | gtaacgccagggttttcccagtcacgacgaatatt gtctatgcaactccgaaggcggcgcgg | gDNA of *U. maydis* | flanking regions for deletion |
| 22 | MG109_mtf1_LBrv | caattgtcacgccatggtggccatctaggccggc gttgagccaagtagcgaactacccagggg | gDNA of *U. maydis* | flanking regions for deletion |
| 23 | MG110_mtf1_RBfw | gtgcggccgcattaataggcctgagtggccgtg atagattcgattgaccgtattctgtaatg | gDNA of *U. maydis* | flanking regions for deletion |
| 24 | MG111_mtf1_RBrv | gcggataacaatttcacacaggaaacagcaatat tccagaagcgaccgaccgcacatgttgctc | gDNA of *U. maydis* | flanking regions for deletion |
| 25 | MH420_aox1_LBfw | gtaacgccagggttttcccagtcacgacgaatatt cgacacaaacctcaacgtgttcaacgttg | gDNA of *U. maydis* | flanking regions for deletion |
| 26 | MH421_aox1_LBrv | caattgtcacgccatggtggccatctaggcccctt gtacacagtggtggtcgagcgtaaaaag | gDNA of *U. maydis* | flanking regions for deletion |
| 27 | MH422_aox1_RBfw | gtgcggccgcattaataggcctgagtggccaaa cacaccaggtacaaagcacgtgatcctgtaaac | gDNA of *U. maydis* | flanking regions for deletion |
| 28 | MH423_aox1_RBrv | gcggataacaatttcacacaggaaacagcaatat tagacactgttgttggtgcagcggaacc | gDNA of *U. maydis* | flanking regions for deletion |
| 29 | MH424_vbs1_LBfw | gtaacgccagggttttcccagtcacgacgaatatt acccacgccaggcgcagagagaccgttgg | gDNA of *U. maydis* | flanking regions for deletion |
| 30 | MH425_vbs1_LBrv | caattgtcacgccatggtggccatctaggcccttg aagaagctcagatgtctaggtcagc | gDNA of *U. maydis* | flanking regions for deletion |

| | | | | |
|---|---|---|---|---|
| 31 | MH426_vbs1_RBfw | gtgcggccgcattaataggcctgagtggccgtc gtggatatgacttgcagttttactctgtac | gDNA of *U. maydis* | flanking regions for deletion |
| 32 | MH427_vbs1_RBrv | gcggataacaatttcacacaggaaacagcaatat tcgccagctgtcaattgtacacacctgg | gDNA of *U. maydis* | flanking regions for deletion |
| 33 | MH428_orf4_LBfw | gtaacgccagggttttcccagtcacgacgaatatt gacactgcttccaccacccaggccaaaac | gDNA of *U. maydis* | flanking regions for deletion |
| 34 | MH429_orf4_LBrv | caattgtcacgccatggtggccatctaggccagt atcgttgagggctgtacaagttcgag | gDNA of *U. maydis* | flanking regions for deletion |
| 35 | MH430_orf4_RBfw | gtgcggccgcattaataggcctgagtggccagc aactcaacgtggtccatttcaatatcaatc | gDNA of *U. maydis* | flanking regions for deletion |
| 36 | MH431_orf4_RBrv | gcggataacaatttcacacaggaaacagcaatat tgcttcgctcatgcctcagacgcgctccac | gDNA of *U. maydis* | flanking regions for deletion |
| 37 | MH432_pks3_LBfw | gtaacgccagggttttcccagtcacgacgaatatt ggataagagcaagaagctgacgcaagctc | gDNA of *U. maydis* | flanking regions for deletion |
| 38 | MH433_pks3_LBrv | caattgtcacgccatggtggccatctaggccgcc gatagattaccaatgtggctaacgctg | gDNA of *U. maydis* | flanking regions for deletion |
| 39 | MH434_pks3_RB fw | gtgcggccgcattaataggcctgagtggccaaa cgatcgagccaatgttgcctgacagacgtc | gDNA of *U. maydis* | flanking regions for deletion |
| 40 | MH435_pks3_RB rv | gcggataacaatttcacacaggaaacagcaatat tctccttactcgaacttgtacagccctc | gDNA of *U. maydis* | flanking regions for deletion |
| 41 | MH436_omt1_LBfw | gtaacgccagggttttcccagtcacgacgaatatt ccgggatctcgtcgatcgcttgagctccc | gDNA of *U. maydis* | flanking regions for deletion |
| 42 | MH437_omt1_LBrv | caattgtcacgccatggtggccatctaggcccct caaagaaagaggctagttttcaggaaaac | gDNA of *U. maydis* | flanking regions for deletion |
| 43 | MH438_omt1_RBfw | gtgcggccgcattaataggcctgagtggccgca cctcttgcaagaacgctatcgagttcttc | gDNA of *U. maydis* | flanking regions for deletion |
| 44 | MH439_omt1_RBrv | gcggataacaatttcacacaggaaacagcaatat tcaggtgacaagcatcaccaagtagtatc | gDNA of *U. maydis* | flanking regions for deletion |
| 45 | MH440_pmo1_LBfw | gtaacgccagggttttcccagtcacgacgaatatt cagctctccaaccgagcagcccatgg | gDNA of *U. maydis* | flanking regions for deletion |
| 46 | MH441_pmo1_LBrv | caattgtcacgccatggtggccatctaggccgttt ttcaagtgccatataatcgagtaaac | gDNA of *U. maydis* | flanking regions for deletion |

| | | | | |
|---|---|---|---|---|
| 47 | MH442_pmo1_RBfw | gtgcggccgcattaataggcctgagtggcccca caatcttgaagcttttccctcacgcctgtc | gDNA of *U. maydis* | flanking regions for deletion |
| 48 | MH443_pmo1_RBrv | gcggataacaatttcacacaggaaacagcaatat tgaaaacaggaatgccgcgaactagatgac | gDNA of *U. maydis* | flanking regions for deletion |
| 49 | MH444_orf5_LBfw | gtaacgccagggttttcccagtcacgacgaatatt ggcagagttgagcaccactatgcggttgc | gDNA of *U. maydis* | flanking regions for deletion |
| 50 | MH445_orf5_LBrv | caattgtcacgccatggtggccatctaggccgttt cgtctgcagaagagctgctgtatac | gDNA of *U. maydis* | flanking regions for deletion |
| 51 | MH446_orf5_RBfw | gtgcggccgcattaataggcctgagtggccaac acgcgcttgatcgactcactcctcagtcctc | gDNA of *U. maydis* | flanking regions for deletion |
| 52 | MH447_orf5_RBrv | gcggataacaatttcacacaggaaacagcaatat ttcttgacccacaccgaccatgcgcagcac | gDNA of *U. maydis* | flanking regions for deletion |
| 53 | MH448_cyp4_LBfw | gtaacgccagggttttcccagtcacgacgaatatt cggatcaacttggtggctagtccagggc | gDNA of *U. maydis* | flanking regions for deletion |
| 54 | MH449_cyp4_LBrv | caattgtcacgccatggtggccatctaggccgtc gatcaacaaatacacgtccacagcg | gDNA of *U. maydis* | flanking regions for deletion |
| 55 | MH450_cyp4_RBfw | gtgcggccgcattaataggcctgagtggccacc acacatctttctgcaataggcaatgcg | gDNA of *U. maydis* | flanking regions for deletion |
| 56 | MH451_cyp4_RBrv | gcggataacaatttcacacaggaaacagcaatat tctcgtcacaggcagcaccgatcctgaagc | gDNA of *U. maydis* | flanking regions for deletion |
| 57 | MH452_deh1_LB fw | gtaacgccagggttttcccagtcacgacgaatatt gcgatcgacgctccaatcagcggtggtgc | gDNA of *U. maydis* | flanking regions for deletion |
| 58 | MH453_deh1_LBrv | caattgtcacgccatggtggccatctaggccttcg tacagcttgcaagatcgagaaagcg | gDNA of *U. maydis* | flanking regions for deletion |
| 59 | MH454_deh1_RB fw | gtgcggccgcattaataggcctgagtggccaatt cagtgcgacgctatccgtttgaatcgcc | gDNA of *U. maydis* | flanking regions for deletion |
| 60 | MH455_deh1_RB rv | gcggataacaatttcacacaggaaacagcaatat tgacggctctggactcaccgccgagggag | gDNA of *U. maydis* | flanking regions for deletion |
| 61 | MG700_mtf1_XmaIfw | gcatcccgggccatggctggaaaacgtaatcgc | gDNA of *U. maydis* | overexpression of *mtf1* in *U. maydis* |
| 62 | MG703_mtf1_NotIrv | atggcggccgctcagaccacggtgttagtggc | gDNA of *U. maydis* | overexpression of *mtf1* in *U. maydis* |

| | | | | |
|---|---|---|---|---|
| 63 | MG554_mtf2_XmaIfw | gcatcccgggatgtcttgcccacaggatcgc | gDNA of *U. maydis* | overexpression of *mtf2* in *U. maydis* |
| 64 | MG551_mtf2_NotIrv | atgcgcggccgcctagtgaaaagggcgttggcg | gDNA of *U. maydis* | overexpression of *mtf2* in *U. maydis* |
| 65 | MH701_pks3_XmaIfw | gcatcccgggccatgtcaagtcaaagttttgc | gDNA of *U. maydis* | overexpression of *pks3* in *U. maydis* |
| 66 | MH702_pks3_NotIrv | atggcggccgcctactgtgatagtggcttcg | gDNA of *U. maydis* | overexpression of *pks3* in *U. maydis* |
| 67 | MI985_crg_SbfIfw | atatcctgcaggctgggaccataccgtgttgc | plasmid | overexpression of *pks3* in *U. maydis* |
| 68 | MI986_pks3_AflIIrv | atatcttaagaaactttattgccaaatgtttg | plasmid | overexpression of *pks3* in *U. maydis* |
| 69 | MI593_pks4_XmaIfw | gcatcccgggccatgtcttcccactcctctagtc | gDNA of *U. maydis* | overexpression of *pks4* in *U. maydis* |
| 70 | MH704_pks4_NotIrv | cggatgagctgggatcactatag | gDNA of *U. maydis* | overexpression of *pks4* in *U. maydis* |
| 71 | MI059_cyp4_XmaIfw | gcatcccgggccatgttcgctctcgaggtagatg | gDNA of *U. maydis/*plasmid | overexpression of *cyp4* in *U. maydis/*probe for Northern blot |
| 72 | MI060_cyp4_NotIrv | atggcggccgctcaatccttgtagtagtgatcg | gDNA of *U. maydis/*plasmid | overexpression of *cyp4* in *U. maydis/*probe for Northern blot |
| 73 | MH86_pks5_fw | cgagcacggtgttgagctcttg | gDNA of *U. maydis* | probe for Northern blot |
| 74 | MH87_pks5_rv | cgcttgtaggataacaccgc | gDNA of *U. maydis* | probe for Northern blot |
| 75 | MH88_orf1_fw | cgactttccgttttggtgagtagc | gDNA of *U. maydis* | probe for Northern blot |
| 76 | MH89_orf1_rv | gcttcaatggccaatactgcacg | gDNA of *U. maydis* | probe for Northern blot |
| 77 | MH90_pks4_fw | cgtttcgccactgctatcgatcaagc | gDNA of *U. maydis* | probe for Northern blot |
| 78 | MH91_pks4_rv | cgtaagcttggcgtccgtccaggc | gDNA of *U. maydis* | probe for Northern blot |
| 79 | MH92_vbs1_fw | gctgtttctccatatgatgc | gDNA of *U. maydis* | probe for Northern blot |
| 80 | MH93_vbs1_rv | ggatcacagtcacaagctgc | gDNA of *U. maydis* | probe for Northern blot |
| 81 | MH94_orf4_fw | gctgctcagacgcaaaaccc | gDNA of *U. maydis* | probe for Northern blot |
| 82 | MH95_orf4_rv | gcatcaatgagtactggatgg | gDNA of *U. maydis* | probe for Northern blot |
| 83 | MH96_pks3_fw | cggtagggcggaaggcgttgc | gDNA of *U. maydis* | probe for Northern blot |

| | | | | |
|---|---|---|---|---|
| 84 | MH97_pks3_rv | gctggaaatgtccaggtgtgtgg | gDNA of *U. maydis* | probe for Northern blot |
| 85 | MH98_omt1_fw | gctctcctcgagaattaccagc | gDNA of *U. maydis* | probe for Northern blot |
| 86 | MH99_omt1_rv | cgaaccttggccgagtgcagg | gDNA of *U. maydis* | probe for Northern blot |
| 87 | MH100_pmo1_fw | cgaaccttctgggatacaagcc | gDNA of *U. maydis* | probe for Northern blot |
| 88 | MH101_pmo1_rv | gcattctcctgctcggtgttgc | gDNA of *U. maydis* | probe for Northern blot |
| 89 | MH102_orf5_fw | cgtcagcgacacctgatctgc | gDNA of *U. maydis* | probe for Northern blot |
| 90 | MH103_orf5_rv | cgtgaccctatcgaggccatgacc | gDNA of *U. maydis* | probe for Northern blot |
| 91 | MH106_deh1_fw | cgttctcgtcacaggcagcacc | gDNA of *U. maydis* | probe for Northern blot |
| 92 | MH107_deh1_rv | cgatggaacctcgagcagagg | gDNA of *U. maydis* | probe for Northern blot |
| 93 | MH108_orf6_fw | cgtcagatggcgggctgggactcg | gDNA of *U. maydis* | probe for Northern blot |
| 94 | MH109_orf6_rv | cgacacgcgatgcgacaccgatgg | gDNA of *U. maydis* | probe for Northern blot |
| 95 | MH110_orf7_fw | gcaacgtcaccaaagccttgacc | gDNA of *U. maydis* | probe for Northern blot |
| 96 | MH111_orf7_rv | ggtgtactcgcctgcgtacgacg | gDNA of *U. maydis* | probe for Northern blot |
| 97 | MG1012_mtf2_fw | cgttaaacgtatcgaagctagc | gDNA of *U. maydis* | probe for Northern blot |
| 98 | MG1013_mtf2_rv | gcaacacaagatcgtgaggagc | gDNA of *U. maydis* | probe for Northern blot |
| 99 | MG1014_orf2_fw | atggatcagcacaagcgaggc | gDNA of *U. maydis* | probe for Northern blot |
| 100 | MG1015_orf2_rv | ttagaacaagatgagaacgtgtctcctgc | gDNA of *U. maydis* | probe for Northern blot |
| 101 | MG1016_orf3_fw | atgagaagcgcagcaatcgaagc | gDNA of *U. maydis* | probe for Northern blot |
| 102 | MG1017_orf3_rv | tcatccatgtggacaagtggc | gDNA of *U. maydis* | probe for Northern blot |
| 103 | MG1018_mtf1_fw | gctcaaaaaatctgaatggaccg | gDNA of *U. maydis* | probe for Northern blot |
| 104 | MG1019_mtf1_rv | cgtgccgggtcgccagccatgg | gDNA of *U. maydis* | probe for Northern blot |
| 105 | MG1020_aox1_fw | cgacgaatcaaacaggtctacactgg | gDNA of *U. maydis* | probe for Northern blot |
| 106 | MG1021_aox1_rv | cgaagcgggcgcgatgcgatcg | gDNA of *U. maydis* | probe for Northern blot |
| 107 | ME48_pRS426_fw | gggggatgtgctgcaaggcg | plasmid | sequencing |
| 108 | ME49_pRS426_rv | tccggctcctatgttgtgtgg | plasmid | sequencing |
| 109 | MH758_pks3_fw | cactttctcagcccagacggaaag | plasmid | sequencing |
| 110 | MH759_pks3_fw | cgactcttagagccagagaaaatgc | plasmid | sequencing |
| 111 | MH760_pks3_fw | cgaaatagacaccgacttgcag | plasmid | sequencing |
| 112 | MH761_pks3_fw | gccagctgaaacgaatcactgag | plasmid | sequencing |

| | | | | |
|---|---|---|---|---|
| 113 | MH762_pks3_fw | gtcgctcccatgcctgcagagg | plasmid | sequencing |
| 114 | MH763_pks3_fw | gtacttgagcgctgtttgcccg | plasmid | sequencing |
| 115 | MH764_pks3_fw | ggacaaggtcgatcgtctttctg | plasmid | sequencing |
| 116 | MI917_np_pks3_fw | cctctttctttgaggatggcttc | plasmid | sequencing |
| 117 | MI918_np_pks3_fw | cgtccagcgcgatttcacaaatc | plasmid | sequencing |
| 118 | MI919_np_pks3_fw | gctcttttggcgacactggtag | plasmid | sequencing |
| 119 | MI920_np_pks3_fw | gtcgataatcccggccgcgctcttg | plasmid | sequencing |
| 120 | MI952_np_pks3_rv | gactggctcactcggctcatagtg | plasmid | sequencing |
| 121 | MI594_pks4_fw | cggatgagctgggatcactatag | plasmid | sequencing |
| 122 | MI595_pks4_fw | cccactcctctagtccgagctc | plasmid | sequencing |
| 123 | MI596_pks4_fw | gttttgccacctcttcttgcac | plasmid | sequencing |
| 124 | MI597_pks4_fw | cacctatccgactccattatcagc | plasmid | sequencing |
| 125 | MI598_pks4_fw | gtgccacaccctcacaggtcaacg | plasmid | sequencing |
| 126 | MI950_pks4_fw | ccaaccgacctcgtgctctctg | plasmid | sequencing |
| 127 | MI120_pks5_fw | cctggctttgtggcaacgcggcg | plasmid | sequencing |
| 128 | MI121_pks5_fw | gggagctgatgctggctttgtattg | plasmid | sequencing |
| 129 | MI122_pks5_fw | caaatgctggctgaatgccgtgc | plasmid | sequencing |
| 130 | MI123_pks5_fw | cgtatcgacgacattatcgttctag | plasmid | sequencing |
| 131 | MI124_pks5_fw | gcaactgccttcgtgtcgcactatg | plasmid | sequencing |
| 132 | MI125_pks5_fw | ccgacctaagtcaggcggaatgg | plasmid | sequencing |
| 133 | MI126_pks5_fw | ccagctgcctagacagtgttgtcg | plasmid | sequencing |
| 134 | MI127_pks5_fw | gctcgatctggccaaagccaggtc | plasmid | sequencing |
| 135 | MI196_cyp4_fw | cgctctcgaggtagatgagagaagc | plasmid | sequencing |
| 136 | MI197_cyp4_fw | gcaccaagagtcagggctacaag | plasmid | sequencing |
| 137 | MI198_cyp4_fw | ggactcaccgccgagggagagatg | plasmid | sequencing |
| 138 | MI199_cyp4_fw | gccagggcatgcacattgccg | plasmid | sequencing |
| 139 | MI951_cyp4_rv | caggaacttgggatgctgacgtg | plasmid | sequencing |
| 140 | MJ049_np_cyp4_fw | gtattcgagctggacgagcgac | plasmid | sequencing |
| 141 | MJ050_np_cyp4_fw | cggatggtgcaatagcacttctc | plasmid | sequencing |

| | | | | |
|---|---|---|---|---|
| 142 | MJ051_np_cyp4_fw | cctcacatcaccacaacacgag | plasmid | sequencing |
| 143 | MI436 vbs1 fw | caagcatctttggcgcatgctgttg | plasmid | sequencing |
| 144 | MI437_vbs1_fw | gcaagcgtttacgtcaacacg | plasmid | sequencing |
| 145 | MI438_vbs1_fw | gctatctggtattggaccagctc | plasmid | sequencing |
| 146 | MI439_vbs1_fw | cgagtcacgaggaagcgttcgac | plasmid | sequencing |
| 147 | MI440_vbs1_fw | ccgctactgcgtgcacactg | plasmid | sequencing |
| 148 | MI930_crg1_fw | cgtttttttcatgttgcatctcg | plasmid | sequencing |
| 149 | MI931_crg1_fw | cattccaatcagtcacgagtgcac | plasmid | sequencing |
| 150 | MI932_crg1_fw | gctactaactgtctttcgcatctc | plasmid | sequencing |
| 151 | MI933_crg1_fw | gcgtgggctcggatatcggtgg | plasmid | sequencing |
| 152 | MI924_crg1_fw | cgtaaggtggacaccttactg | plasmid | sequencing |
| 153 | MI925_crg1_fw | gctgactcctgtcatgggcaatg | plasmid | sequencing |
| 154 | MI926_crg1_fw | cagccgtgcacgaagatctccgg | plasmid | sequencing |
| 155 | MI955_mtf1_fw | cgaaatgaagatgaaattcttaag | plasmid | sequencing |
| 156 | MI956_mtf1_fw | gttcgtcacctatgtccgcgtcg | plasmid | sequencing |
| 157 | MI957_mtf1_fw | cgaagactctggtcgcctcacc | plasmid | sequencing |
| 158 | MI958_mtf1_fw | ccatgttgcgcttgcccaatttttc | plasmid | sequencing |
| 159 | MI959_mtf1_fw | gcgacaagtgcccagtgtagtcc | plasmid | sequencing |
| 160 | MH594_mtf2_fw | cgacgcctaatcacgacgctcc | plasmid | sequencing |
| 161 | MH595_mtf2_fw | cgcacctacgccacattcaacac | plasmid | sequencing |
| 162 | MH596_mtf2_fw | catggcagatgctcgatcctgg | plasmid | sequencing |
| 163 | MI128_polyN_pks4_fw | ggattggaagcatgtttgcgaattg | plasmid | sequencing |
| 164 | MI129_polyN_pks4_rv | cctggtcgtcaaagcttgtgggc | plasmid | sequencing |
