## Supplementary material for "An Unconventional Melanin Biosynthetic Pathway in *Ustilago maydis*": Table S2

**Table S2. Media used in this study.**

| | |
|---|---|
| **Media used for *U. maydis*** | |
| $YEPS_{light}$ | 10 g/l yeast-extract, 4 g/l peptone, 4 g/l saccharose. |
| YNB | 1.7 g/l yeast nitrogen base, 0.2% (w/v) $(NH_4)_2SO_4$, 5% (w/v) glucose or arabinose; pH 5.6-5.8. |
| PDB-Agar | 24 g/l potato dextrose broth, 15 g/l agar; pH 5.6-5.8. |
| Regeneration-Agar (Reg-Agar) | 10 g/l yeast-extract, 20 g/l peptone, 2% (w/v) saccharose, 182.2 g/l sorbitol, 13 g/l agar. |
| **Media used for *S. cerevisiae*** | |
| YPD | 10 g/l yeast-extract, 20 g/l peptone, 2% (w/v) glucose. |
| Synthetic Complete Medium (SC-Medium) | 1.7 g/l yeast nitrogen base without $(NH_4)_2SO_4$, dropout-mix (depending on the selection marker is additionally added: 0.2 g L-histidine, 0.1 g adenine, 0.2 g leucine, 0.2 g tryptophan, 0.15 g methionine or 0.2 g uracil), 2% (w/v) glucose, 2% (w/v) agar; pH 5.6. |
| Dropout-Mix (-His,-Leu,-Ade,-Trp,-Ura,-Met) | 2.0 g alanine, 2.0 g arginine, 2.0 g aspartic acid, 2.0 g asparagine, 2.0 g cysteine, 2.0 g glutamic acid, 2.0 g glutamine, 2.0 g glycine, 2.0 g inositol, 2.0 g isoleucine, 2.0 g lysine, 2.0 g para-aminobenzoic acid, 2.0 g phenylalanine, 2.0 g proline, 2.0 g serine, 2.0 g threonine, 2.0 g tyrosine, 2.0 g valine. |
| **Media used for *E. coli*** | |
| dYT | 16 g/l tryptone, 10 g/l yeast-extract, 5 g/l NaCl, [agar: 2% (w/v)]. |
