## Supplementary material for "An Unconventional Melanin Biosynthetic Pathway in *Ustilago maydis*": Table S3

**Table S3. Plasmids generated in this study.**

| **Plasmid** | **Purpose** | **Resistance**[1] | **Reference** |
|---|---|---|---|
| pJET1.2 | cloning | - | Fermentas |
| pRS426 | drag and drop | - | Sikorski and Hieter, 1989 |
| pETEF-GFP-Ala6-MMXN | gene overexpression | C | Lab collection |
| pCRG-GFP-Ala6-MMXN | gene overexpression | C | Lab collection |
| pJA2880 | gene overexpression | G | Provided by J. Ast |
| pMM40 | gene overexpression | C | Provided by M. Moretti |
| pMM69 | gene overexpression | G | Provided by M. Moretti |
| pMF1-h | resistance cassette | H | Brachmann *et al.*, 2004 |
| pMF1-g | resistance cassette | G | Brachmann *et al.*, 2004 |
| pDL64 | gene overexpression | C | Provided by D. Lanver |
| pRS426-Δ*pks5* | gene deletion | G | this study |
| pRS426-Δ*orf1* | gene deletion | G | this study |
| pRS426-Δ*pks4* | gene deletion | H/G | this study |
| pRS426-Δ*mtf2* | gene deletion | H/G | this study |
| pRS426-Δ*orf3* | gene deletion | H | this study |
| pRS426-Δ*mtf1* | gene deletion | H | this study |
| pRS426-Δ*aox1* | gene deletion | H | this study |
| pRS426-Δ*vbs1* | gene deletion | H | this study |
| pRS426-Δ*orf4* | gene deletion | H | this study |
| pRS426-Δ*pks3* | gene deletion | H | this study |
| pRS426-Δ*omt1* | gene deletion | H | this study |
| pRS426-Δ*pmo1* | gene deletion | H | this study |
| pRS426-Δ*orf5* | gene deletion | H | this study |
| pRS426-Δ*cyp4* | gene deletion | H | this study |
| pRS426-Δ*deh1* | gene deletion | H | this study |
| pCRG-Mtf1-Tnos-Cbx | gene overexpression | C | this study |
| pCRG-Mtf2-Tnos-Cbx | gene overexpression | C | this study |
| pCRG-Pks3-Tnos-Cbx | gene overexpression | C | this study |
| pCRG-Pks3-Tnos-Cbx-G418 | gene overexpression | G | this study |
| pCRG-Pks4-Tnos-Cbx | gene overexpression | C | this study |

[1] Hygromycin resistance (H), Carboxin resistance (C), Geneticin resistance (G)
