## Supplementary material for "An Unconventional Melanin Biosynthetic Pathway in *Ustilago maydis*": Table S4

**Table S4. Strains used in this study.**

| Strain | Genotype | Resistance[1] | Reference |
|---|---|---|---|
| FB1 | *a1 b1* | - | Banuett and Herskowitz, 1989 |
| MB215 | *a2 b13* | - | Lab collection (Hewald) |
| P*crg*::*mtf1* | *a2 b13* P*crg::UMAG_04101* | C | this study |
| P*crg*::*mtf1* Δ*pks5* | *a1b1* P*crg::UMAG_04101* Δ*UMAG_04095* | C, G | this study |
| P*crg*::*mtf1* Δ*orf1* | *a1b1* P*crg::UMAG_04101* Δ*UMAG_04096* | C, G | this study |
| P*crg*::*mtf1* Δ*pks4* | *a1b1* P*crg::UMAG_04101* Δ*UMAG_04097* | C, H | this study |
| P*crg*::*mtf1* Δ*pks4** | *a2b13* P*crg::UMAG_04101* ΔΔ*UMAG_04097* | H, C, G | this study |
| P*crg*::*mtf1* Δ*mtf2* | *a2b13* P*crg::UMAG_04101* Δ*UMAG_11110* | C, G | this study |
| P*crg*::*mtf1* Δ*orf3* | *a2b13* P*crg::UMAG_04101* Δ*UMAG_04100* | C, H | this study |
| P*crg*::*mtf1* Δ*aox1* | *a2b13* P*crg::UMAG_04101* Δ*UMAG_11111* | C, H | this study |
| P*crg*::*mtf1* Δ*vbs1* | *a2b13* P*crg::UMAG_04101* Δ*UMAG_11112* | C, H | this study |
| P*crg*::*mtf1* Δ*orf4* | *a2b13* P*crg::UMAG_04101* Δ*UMAG_04104* | C, H | this study |
| P*crg*::*mtf1* Δ*pks3* | *a2b13* P*crg::UMAG_04101* Δ*UMAG_04105* | C, H | this study |
| P*crg*::*mtf1* Δ*omt1* | *a2b13* P*crg::UMAG_04101* Δ*UMAG_04106* | C, H | this study |
| P*crg*::*mtf1* Δ*pmo1* | *a2b13* P*crg::UMAG_04101* Δ*UMAG_04107* | C, H | this study |
| P*crg*::*mtf1* Δ*orf5* | *a2b13* P*crg::UMAG_04101* Δ*UMAG_12253* | C, H | this study |
| P*crg*::*mtf1* Δ*cyp4* | *a2b13* P*crg::UMAG_04101* Δ*UMAG_04109* | C, H | this study |
| P*crg*::*mtf1* Δ*deh1* | *a2b13* P*crg::UMAG_04101* Δ*UMAG_11113* | C, H | this study |
| P*crg*::*mtf1* Δ*cyp4*Δ*vbs1* | *a2b13* P*crg::UMAG_04101* Δ*UMAG_04109* Δ*UMAG_11112* | H, C, G | this study |
| P*crg*::*mtf2* | *a2b13* P*crg::UMAG_11110* | C | this study |
| P*crg*::*pks3* | *a2b13* P*crg::UMAG_04105* | C | this study |
| P*crg*::*pks4* | *a2b13* P*crg::UMAG_04097* | C | this study |
| P*crg*::*pks3* + P*crg*::*pks4* | *a2b13* P*crg::UMAG_04105* + P*crg::UMAG_04097* | C, G | this study |

[1] Hygromycin resistance (H), Carboxin resistance (C), Geneticin resistance (G)
