## Supplementary material for "An Unconventional Melanin Biosynthetic Pathway in *Ustilago maydis*": Table S5

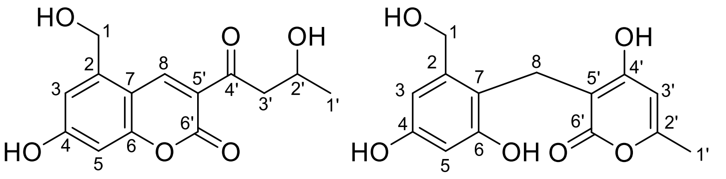

6 7

**Table S5.** **$^{1}$H (500 MHz) and $^{13}$C (125 MHz) NMR spectroscopic data for compounds 6 and 7 in MeOH-$d_4$ ($\delta$ in ppm and $J$ in Hz).**

| No. | 1 | | | 2 | |
|---|---|---|---|---|---|
| | $\delta_H$ (mult., $J$) | $\delta_C$ | | $\delta_H$ (mult., $J$) | $\delta_C$ |
| 1 | 4.87 (s) | 60.7 | | 4.70 (s) | 62.9 |
| 2 | - | 143.8 | | - | 141.1 |
| 3 | 6.91 (d, 2.1) | 114.1 | | 6.34 (d, 2.6) | 107.9 |
| 4 | - | 166.0 | | - | 155.8 |
| 5 | 6.63 (d, 2.1) | 101.0 | | 6.23 (d, 2.6) | 102.9 |
| 6 | - | 158.9 | | - | 157.1 |
| 7 | - | 108.8 | | - | 117.2 |
| 8 | 8.84 (s) | 145.1 | | 3.67 (s) | 19.3 |
| 1' | 1.27 (d, 6.3) | 22.1 | | 2.14 (s) | 17.9 |
| 2' | 4.34 (m) | 63.8 | | - | 159.3 |
| 3' | 3.23 (d, 6.2) | 50.7 | | n.d. | 105.9 |
| 4' | - | 196.6 | | - | 177.2 |
| 5' | - | 117.8 | | - | 99.7 |
| 6' | - | 160.2 | | - | 169.5 |

n.d. = not detected
