## Supplementary material for "An Unconventional Melanin Biosynthetic Pathway in *Ustilago maydis*": Table S6

**Table S6. HR-ESI-MS data of all compounds described in this work.**

| Compounds | Detected mass $[M + H]^+$ | Calculated mass $[M + H]^+$ | Error (ppm) | Sum formula $[M + H]^+$ |
|---|---|---|---|---|
| **1** | 169.0497 | 169.0495 | -1.1 | $C_8H_9O_4$ |
| **2** | 153.0545 | 153.0546 | 0.1 | $C_8H_9O_4$ |
| **3** | 127.0380 | 127.0390 | 1.0 | $C_6H_7O_3$ |
| **4/5** | 263.0913 | 263.0914 | 0.4 | $C_{14}H_{15}O_5$ |
| **6/7** | 279.0864 | 279.0863 | -0.3 | $C_{14}H_{15}O_6$ |
